# Universality of Form: The Case of Retinal Cone Photoreceptor Mosaics

**DOI:** 10.1101/2022.01.09.475540

**Authors:** Alireza Beygi

## Abstract

Cone photoreceptor cells are wavelength-sensitive neurons in the retinas of vertebrate eyes and are responsible for color vision. The spatial distribution of these nerve cells is commonly referred to as cone photoreceptor mosaic. By applying the principle of maximum entropy, we demonstrate the universality of retinal cone mosaics in vertebrate eyes by examining various species, namely, rodent, dog, monkey, human, fish, and bird. We introduce a parameter called retinal temperature, which is conserved across the retinas of vertebrates. The virial equation of state for two-dimensional cellular networks, known as Lemaître’s law, is also obtained as a special case of our formalism. We investigate the behavior of several artificially generated networks and the natural one of the retina concerning this universal, topological law.

## 1. Introduction

The principle of maximum entropy provides an estimation for the underlying probability distribution of the observed data that corresponds best to the currently available information about the system [1,2]. It has been applied to fields as diverse as physics [3,4], biology [5], ecology [4,6], and natural language [7]. The philosophy behind the maximum entropy inference approach is to explain and predict experimental observations by making the fewest number of assumptions (i.e., constraints) while assuming no explicit underlying mechanisms.

One of the prime and dreadfully arduous challenges in applying the principle of maximum entropy to a given system is to find out relevant constraints that should be imposed on the system [8]. The authors of [9] have suggested that in situations where the experiments are repeatable, the expected value of the entropy of the likelihood function is relevant information that should be considered a constraint. Although, for a given system, its value is largely unknown. Solving the corresponding Lagrange problem leads to the so-called entropic probability distribution [10,11]. Entropic distributions have been exploited mainly within the context of data classification and theoretical physics [12]. Yet the consequences of such an approach are not fully explored in biology and life sciences. In the context of biology, due to the rigid structure of DNA, most experiments must be repeatable.

In the present paper, we adopt the approach of [9] and apply it to a complex multicellular biological system, namely, cone photoreceptor cells in the retina; for an earlier attempt in this direction, see [13]. Cone cells are wavelength-sensitive receptors in the retinas of vertebrate eyes, and their different sensitivities and responses to light of different wavelengths mediate color vision. The spatial distribution of these cells, so-called cone mosaic [14], varies among different species, which in each case may reflect the evolutionary pressures that give rise to various adaptations to the lifestyle of a particular species and its specific visual needs. Although, in most cases, the adaptive value of a particular cone mosaic is unknown [15]. From the perspective of gene regulatory mechanisms, the most fundamental questions such as: what are the mechanisms which control the mostly random distributions of cone subtypes in the human retina? or, what migration mechanisms determine the highly regular and ordered patterns of cone subtypes in the retina of the zebrafish? remain unanswered [16].

In the current work, we show that various forms of distributions of cone cells are controlled by entropy, and we predict the frequency of the appearance of cones in the retina. To this end, we employ the principle of maximum entropy without invoking any specific biological mechanisms or driving forces. In a nutshell, we look for a configuration of sensory cells that maximizes entropy while the expected value of the entropy of the likelihood—which codifies information about the local environment of cells—has been imposed as a constraint. One of the outcomes of this approach is that a configuration with a lower entropy has a higher probability of occurrence (i.e., the frequency of the appearance). This approach enables us to identify a conserved retinal factor, which we call retinal *temperature* or *coldness*, in divergent species of rodent, dog, monkey, human, fish, and bird. To our knowledge, this is the first model capable of predicting the probability of the occurrence of cone cells in various species’ eyes by tuning a single parameter. For earlier entropic approaches to study neuronal mosaics, see [17,18].

The virial equation of state for two-dimensional cellular networks, known as Lemaître’s law [19,20], relates the fraction of hexagons in a given network to the width of the polygon distribution. Here, we demonstrate how by assuming additional information concerning the topology of the network in the entropy maximization procedure, we can obtain this universal law.

The idea that the organization of biological systems stems from an underlying optimization problem goes back to D’Arcy Thompson, which in his seminal work, *On Growth and Form* [21], he argues for the case of energy minimization, which leads to, for example, the prediction of cellular packing geometries in two-dimensional (2D) networks [22]. The geometric properties, obtained based on the knowledge of the physical properties of epithelial cells, can be considered as factors that control the development and function of a living organism [21]. Reducing seemingly different phenomena to a simple governing principle was the manifestation of the universality of form to Thompson [23]. In essence, here, we are replacing energy minimization with entropy maximization, with the advantage of ignoring the involved forces and physical interactions, which incidentally implies a mathematical (entropic) restriction on the evolution of biological forms.

This paper is organized as follows. In Section 2, we review the problem of entropy maximization as applied in this paper. We study the spatial distributions of cone cells in the retinas of various vertebrates in Section 3 and demonstrate the predictive power of our approach besides its explanatory nature. In Section 4, we derive Lemaître’s law and examine it in several artificially generated cellular networks and cone mosaics. We summarize and conclude in Section 5.

## 2. Entropy Maximization

In statistical mechanics, to obtain the Boltzmann distribution from the principle of maximum entropy, one has to assume a constraint on the mean energy value as, in the context of physics, the expected value of energy is crucial information about the system. This approach leads to a formalism in which thermodynamic temperature emerges as a free parameter and should be determined later from experiment [24]. In a general setting, the challenge is to find out relevant constraints that should be imposed on the system. A. Caticha and R. Preuss in [9] have assumed a set of data generated by some experiment, where the only requirement is the experiment to be repeatable. If, for example, the experiment is performed twice, with the corresponding outcomes of *z*_1_ and *z*_2_, in the case that we discard the value of *z*_2_, the resulting situation should be indistinguishable from if we had done the experiment only once. They have argued that a constraint on the expected value of the entropy of the likelihood codifies this information. Inspired by this idea and since biological experiments must be repeatable because of the robustness of DNA, we adopt this specific approach to entropy maximization and apply it to multicellular biological systems; see also [13] and the references cited therein. Generally, an experiment does not need to be repeatable; for instance, this may be the case at the atomic scale [25]. For non-repeatable experiments, the Einstein fluctuation formula is applicable [9,26]. Note that in cellular biology, each cell is composed of a large number of atoms, and the experiments are robust and repeatable.

We denote sensory neurons *S* and their local environment, which consists of other cells *Y*. We assume the following information about the system:

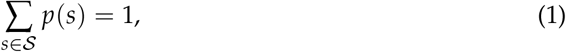

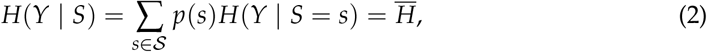

where 𝒮 denotes the support set of *S*. Equation (1) is a normalization condition of the probability mass function (in this paper, the frequency of the appearance) of neurons, Equation (2) assumes the knowledge of the numerical value 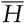 of *H*(*Y* | *S*), and *H*(*Y* | *S* = *s*) = − ∑_*y*∈𝒴_ *f* (*Y* = *y* | *S* = *s*) ln *f* (*Y* = *y* | *S* = *s*), where 𝒴 denotes the support set of *Y*, is defined in terms of the probabilities *f* (*Y* | *S* = *s*). By the method of Lagrange multipliers, we maximize the Shannon entropy of neurons, *H*(*S*) = − ∑_*s*_ ∈ 𝒮*p*(*s*) ln *p*(*s*), while taking (1) and (2) into account. The corresponding Lagrangian reads

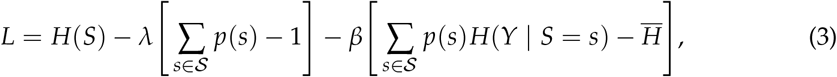

where *λ* and *β* are Lagrange multipliers. By solving *∂L*/*∂p*(*s*) = 0, we obtain the so-called entropic probability [9–11]:

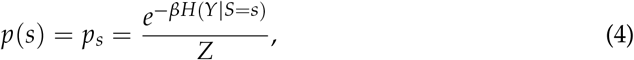

where *Z* = ∑_*w∈*_*𝒮* exp [− *βH*(*Y S* = *w*)]. Assuming *β* > 0, Equation (4) implies that neurons with lower entropy *H*(*Y S* = *s*) have a higher probability or frequency of appearance, which is confirmed in the case of cone photoreceptors in Section 3. The probability distribution in (4) is the most likely and the least-biased one, where the only assumed knowledge about the system is the repeatability nature of the experiments. Other available information about the system can be incorporated as additional constraints in (3); an example of such a scenario is given in Section 4.

A couple of remarks are in order. The application of the principle of maximum entropy strongly depends on how we specify the system configuration, which by itself depends on the nature of the problem at hand. Different ways of describing the configuration of the same system may lead to different outcomes; for a detailed discussion of this issue, see [27]. The second remark deals with (2). Although we have assumed the knowledge of 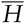, we do not know its value in most cases, but rather, it is a quantity that its value *should* be known, thus we have formulated our problem as if we had this information; for a detailed discussion of this matter, see [9]. By calculating the free parameter, *β*, from the experimental data, one can infer the value of 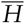. In analogy with statistical mechanics where thermodynamic temperature emerges as the inverse of the Lagrange multiplier in the derivation of the Boltzmann distribution, we interpret *β* as the biological *coldness* (the reciprocal of *temperature*) of neurons. As in thermodynamics, where energy is an extensive quantity, here entropy is also extensive. Note that thermodynamic temperature is a statistical property of matter in bulk, and thus *β* can be viewed as an emergent quantity at a tissue level.

## 3. Spatial Distributions of Cone Photoreceptors in the Retinas of Vertebrates

In this section, we apply the principle of maximum entropy, culminated in Equation (4), to retinal cone mosaics of various vertebrates. Besides predicting the frequency of the appearance of cones across the retina, we demonstrate that the application of the maximum entropy inference leads to the introduction of a new parameter, which we call retinal coldness, that is conserved in divergent species of rodent, dog, monkey, human, fish, and bird. In Section 3.1, we elaborate on the details of our calculations in the case of human cones; other species are summarized in Section 3.2.

### 3.1. Spatial Distributions of Human Cone Photoreceptors

Human color vision is mediated by three types of cones, which are sensitive to (blue) short-, (green) medium-, and (red) long-wavelength light. The spatial distributions of these cells in a living human eye are shown in Figure 1. The image in the top-left corner is the first image of the spatial arrangement of living human cones, reported in [28].

**Figure 1.**
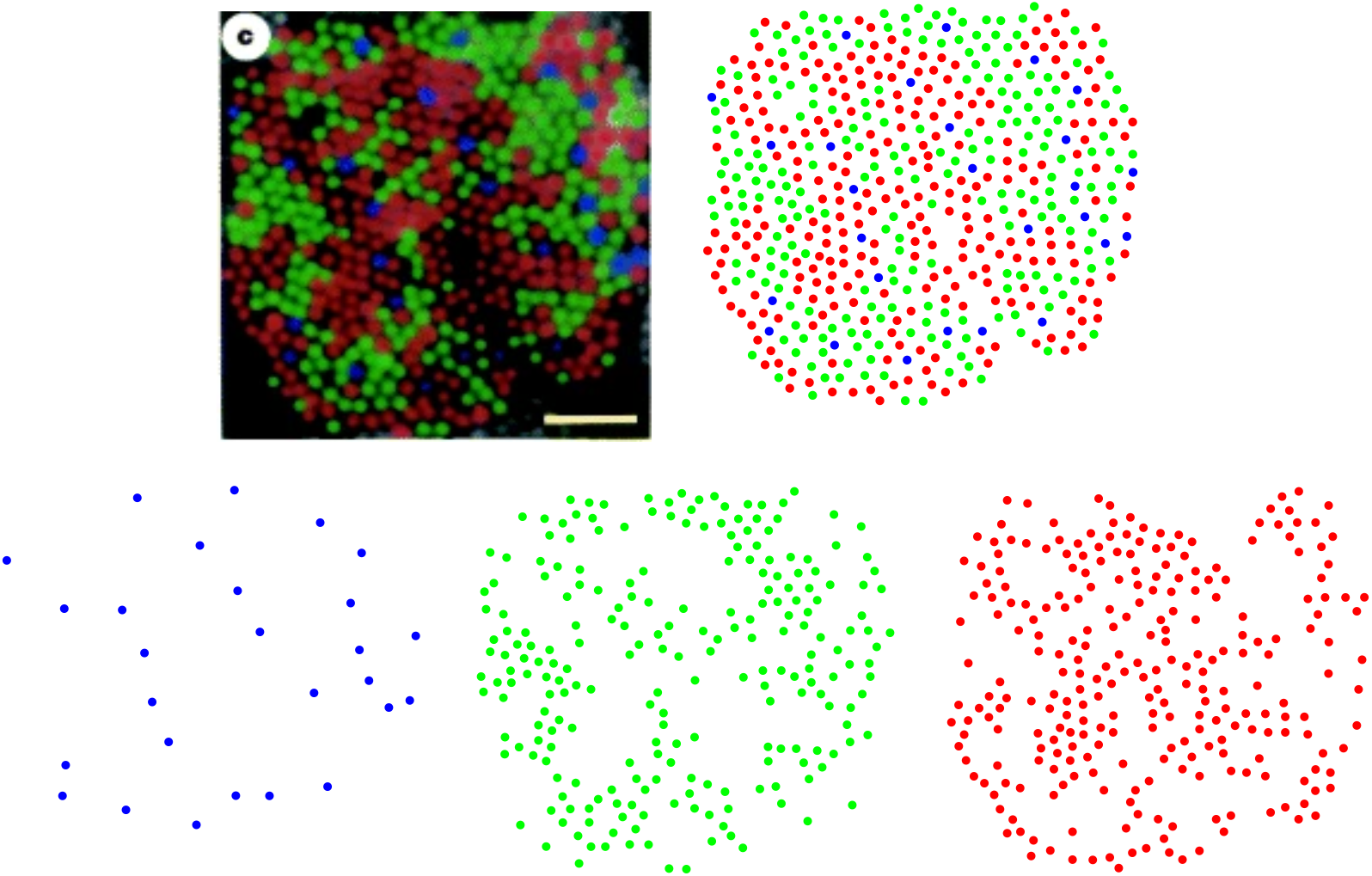
Spatial distributions of cone photoreceptors in a living human nasal retina, at one degree of eccentricity. The image, in the top-left corner, is adapted from [28], where the scale bar = 5 µm. Figures in the bottom row, from left to right, illustrate short-, medium-, and long-wavelength-sensitive cones separately.

In the following, we show how Equation (4) can be used to predict the frequency of the appearance of blue, green, and red cones in a retinal field of a human eye given in Figure 1. From (4), we have:

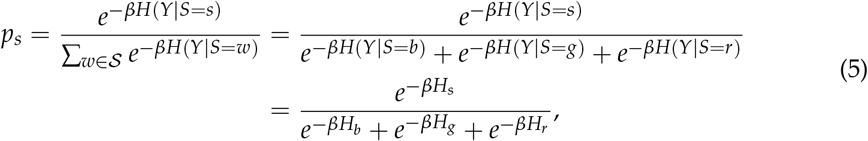

where *p*_*s*_ is the probability of the occurrence or the frequency of the appearance of cone subtypes: blue (*b*), green (*g*), and red (*r*). We consider the local environment of blue cones consists of other blues and exclude green and red cones; the local environment of green cones comprises only greens and the same for the local environment of red cones. This is justified as it is suggested that most cone cells form independent mosaics, and there are no spatial interactions between two mosaics [29]. Also, cone mosaics are explicitly shown to be spatially independent in the case of avian cones [30]. To calculate *H*_*b*_, *H*_*g*_, and *H*_*r*_, we need to consider some kind of probability distribution or density function. Our choice is to construct the nearest-neighbor-distance (NND) distribution for each cone subtype and calculate its corresponding entropy. The rationale behind choosing this specific distribution is as follows. (I) The choice of probability distribution should reflect the frequency with which each cone subtype appears in the retina, which is related to the mean distance between cones of the same type. The scattering of the NND distribution, which is quantified by its entropy, decreases with decreasing the average value of the NND distribution [31] and implies a higher frequency of appearance of cones, based on (5). (II) In general, the methods based on the concept of the nearest-neighbor distance have been extensively used to quantify cone mosaics, see for example [14], which turns out to be a simple but powerful concept to analyze spatial patterns. As an illustration, we have shown searching for the nearest neighbors in the case of blue cones in Figure 2.

**Figure 2.**
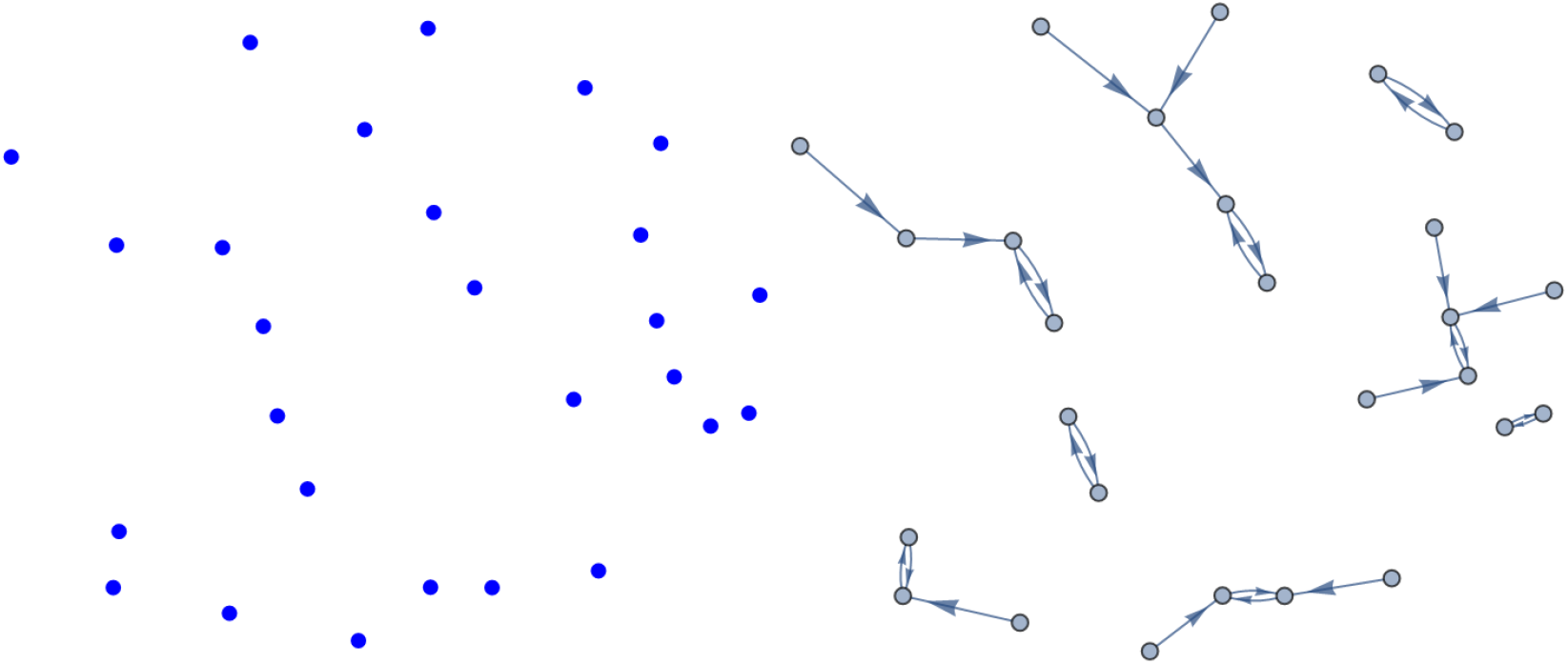
Searching for nearest neighbors, in the case of blue cone photoreceptors in a living human retina.

The NND distribution for each cone subtype is presented in Figure 3. The nearestneighbor distances follow a peaked distribution in each case. Note that to obtain the optimal bin widths of the histograms, we have used a data-based procedure proposed by M. P. Wand [32], to its first-order approximation, which is called one-stage rule (the zeroth-order approximation, i.e., the zero-stage rule, of the method reproduces Scott’s rule of binning [33]).

**Figure 3.**
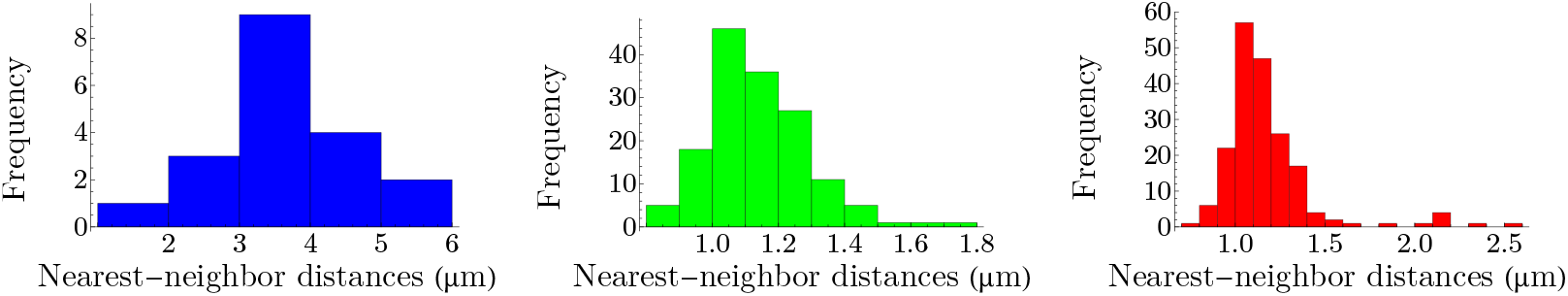
Nearest-neighbor-distance distributions of cone photoreceptors in a living human retina. Values of mean and standard deviation in micrometers for each distribution read *µ*_*b*_ = 3.572, *σ*_*b*_ = 1.020, *µ*_*g*_ = 1.188, *σ*_*g*_ = 0.300, *µ*_*r*_ = 1.172, and *σ*_*r*_ = 0.257.

To calculate entropies of distributions in Figure 3, we use the notion of differential entropy, which is defined as *h*_*s*_ = − ∫ *dx f*_*s*_(*x*) ln *f*_*s*_(*x*), where *f*_*s*_(*x*) is a probability density function, with the property that *dx f*_*s*_(*x*) = 1. We use the notation *H* to designate the Shannon entropy and *h* for the case of differential entropy. Note that as *f*_*s*_(*x*) has units, it cannot be used as the argument of logarithm; however, by the transformation *x* → *x*/*x*_reference_, where *x*_reference_ = 1 µm, we make the distance and subsequently *f*_*s*_(*x*) dimensionless. For each histogram in Figure 3, the density function can be constructed, and the differential entropy can be calculated accordingly; we obtain:

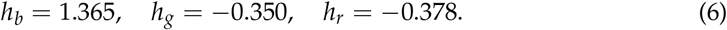

From the image in the top-left corner of Figure 1, we can determine frequencies of appearance of blue, green, and red cones, which are the ratios of these cells in the retinal field. We set the value of *β* in (5) to reproduce the observed values of frequencies. To obtain *β*, we employ the Kullback–Leibler divergence, *D*_KL_ = ∑_*s*_ *q*_*s*_ ln(*q*_*s*_/*p*_*s*_), where *q*_*s*_ corresponds to the observed cone subtype frequency of appearance and *p*_*s*_ corresponds to the prediction of the theory in (5). The left panel of Figure 4 illustrates the Kullback–Leibler divergence as a function of *β*, with the global minimum of 0.001 at *β* = 1.284. The observed cone ratios are compared to the predictions of the theory for *β* = 1.284 in the right panel of Figure 4. Incidentally, these calculations demonstrate that neurons with a lower entropy have a higher probability of occurrence.

**Figure 4.**
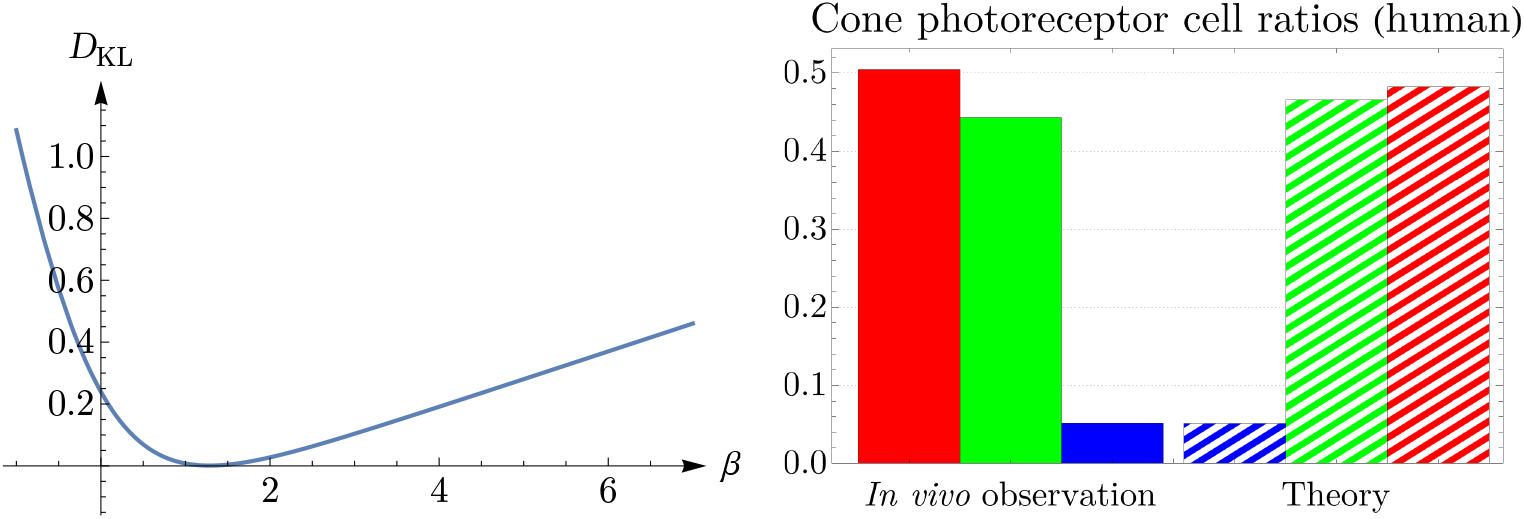
Kullback–Leibler divergence is depicted as a function of *β* in the left panel, where it has the global minimum of 0.001 at *β* = 1.284. The right panel shows a comparison between the *in vivo* observed frequencies of appearance of cone photoreceptors in a human retina and the predictions of the theory (5) for *β* = 1.284.

### 3.2. Spatial Distributions of Vertebrate Cone Photoreceptors: From Rodent to Bird

We apply the procedure explained in Section 3.1 to various vertebrates, namely, rodent, dog, monkey, human, fish, and bird. Rodent and dog are dichromats, monkey, like human, is trichromat, and fish and bird are tetrachromats, which in the case of bird, there is also a significant number of double cones. Our results are summarized in Figures 5, 6, 7, 8, 9, and 10. Although cone mosaics of these diverse species are significantly different from each other, the values of *β* in all species are in the same order, where 1 < *β* < 2. In Section 3.3, by using statistical analyses and the fact that the NND distributions of cone subtypes in vertebrate retinas are peaked—and thus can be approximated by Gaussians—we estimate the value of *β* in a general case.

**Figure 5.**
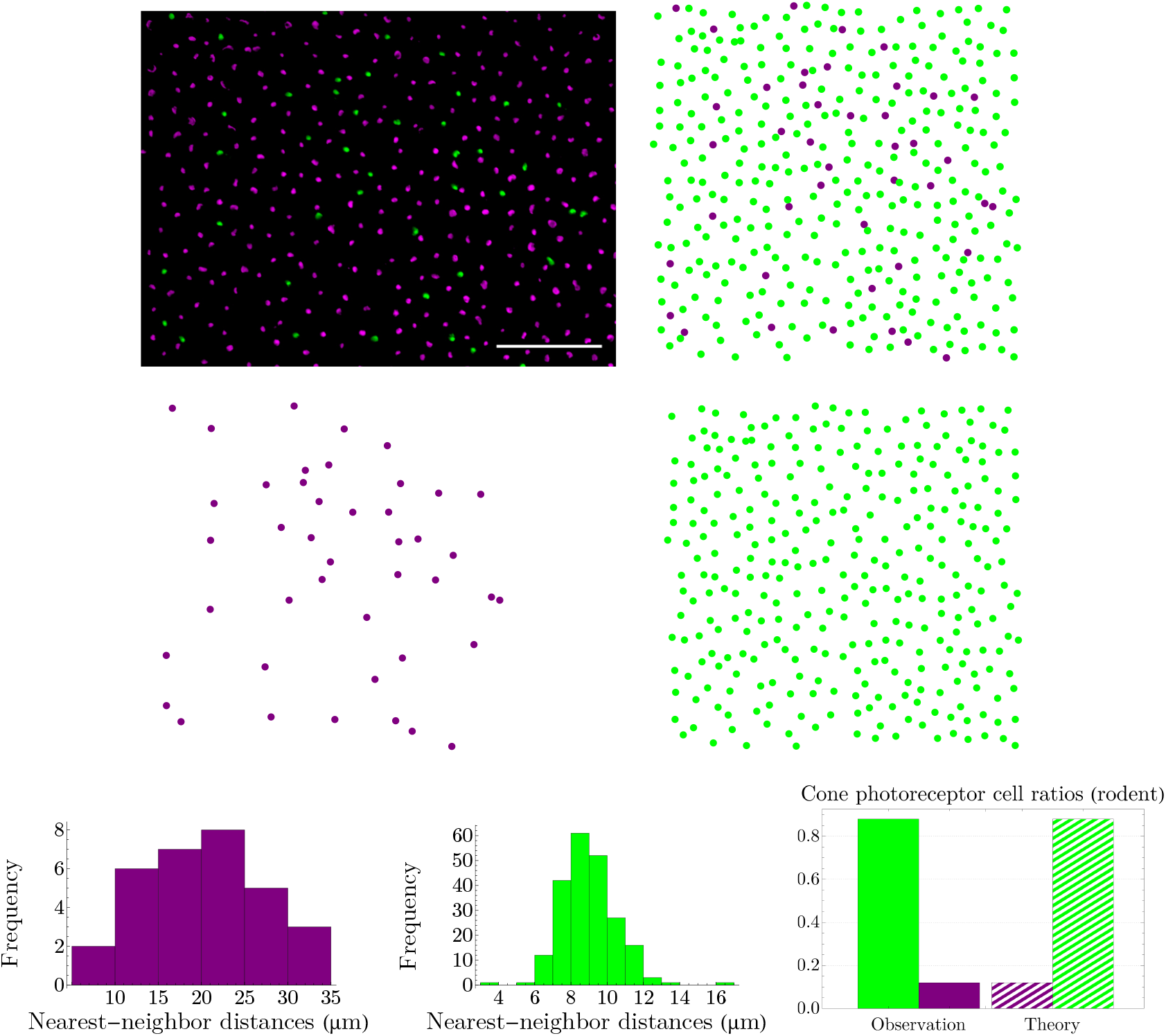
The image in the top-left corner (scale bar = 50 µm), adapted from [34], illustrates the spatial distribution of cone photoreceptors in the dorsal mid-peripheral retina of a diurnal rodent, called the agouti. In this image, the short-wavelength-sensitive-cone opsin is represented as green and the long-wavelength-sensitive-cone opsin as violet; next to it, in the digitized image, we have reversed the colors. Nearest-neighbor-distance distributions in the third row have the entropies of *h*_*v*_ = 3.310 and *h*_*g*_ = 1.787; next to them, we have shown a comparison between the experimental observation of cone ratios in the agouti retina and the predictions of the theory (5) evaluated at the global minimum of the Kullback–Leibler divergence, that is *β* = 1.310.

**Figure 6.**
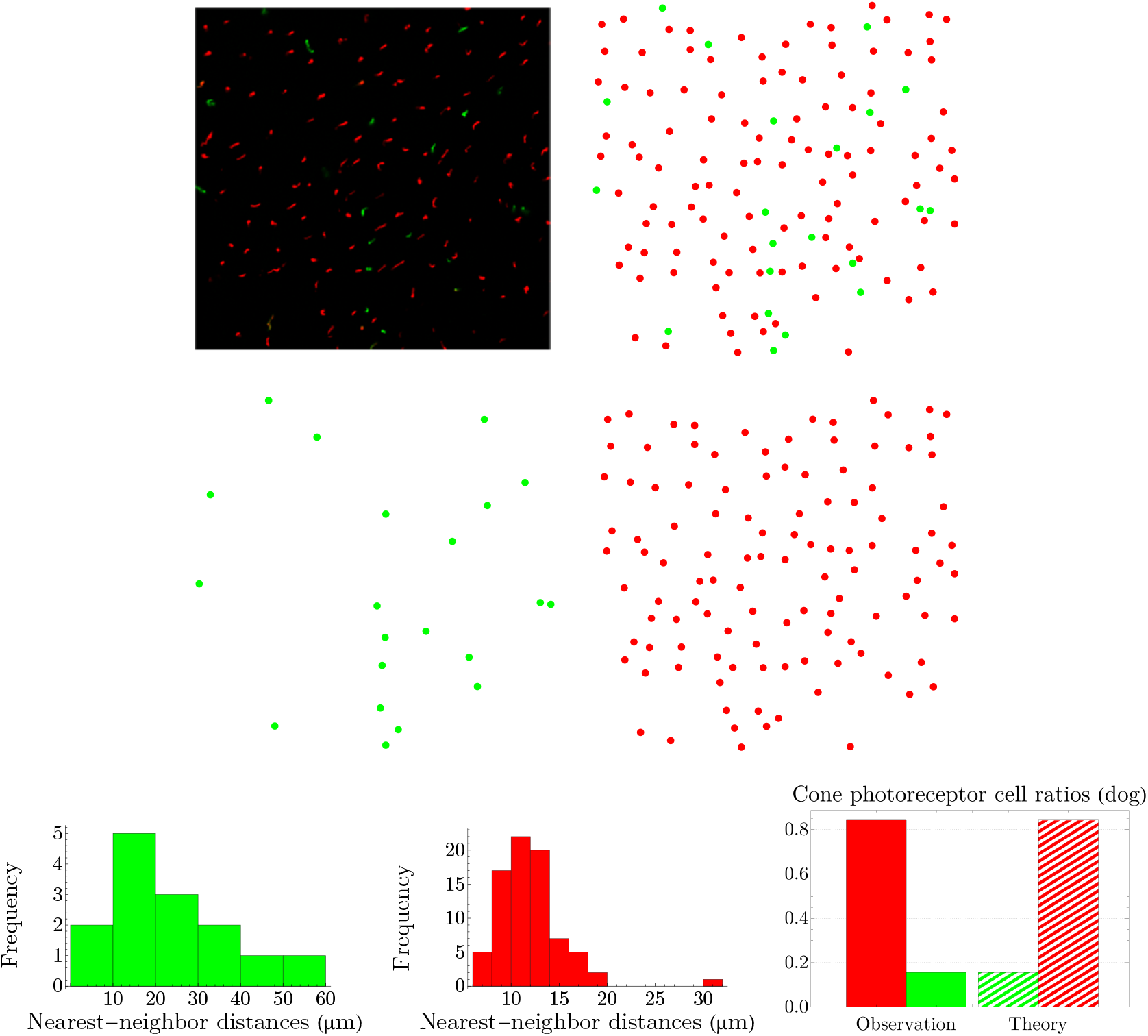
The image in the top-left corner, adapted from [35], shows the spatial distribution of cone photoreceptors in the inferior peripheral retina of a dog; the short-wavelength-sensitive-cone opsin is represented as green, and the long-/medium-wavelength-sensitive-cone opsin as red. The entropies of the NND distributions in the third row read *h*_*g*_ = 3.933 and *h*_*r*_ = 2.440. Next to the NND distributions, we have shown a comparison between the experimental observation of cones’ frequencies of appearance and the predictions of the theory for *β* = 1.127.

**Figure 7.**
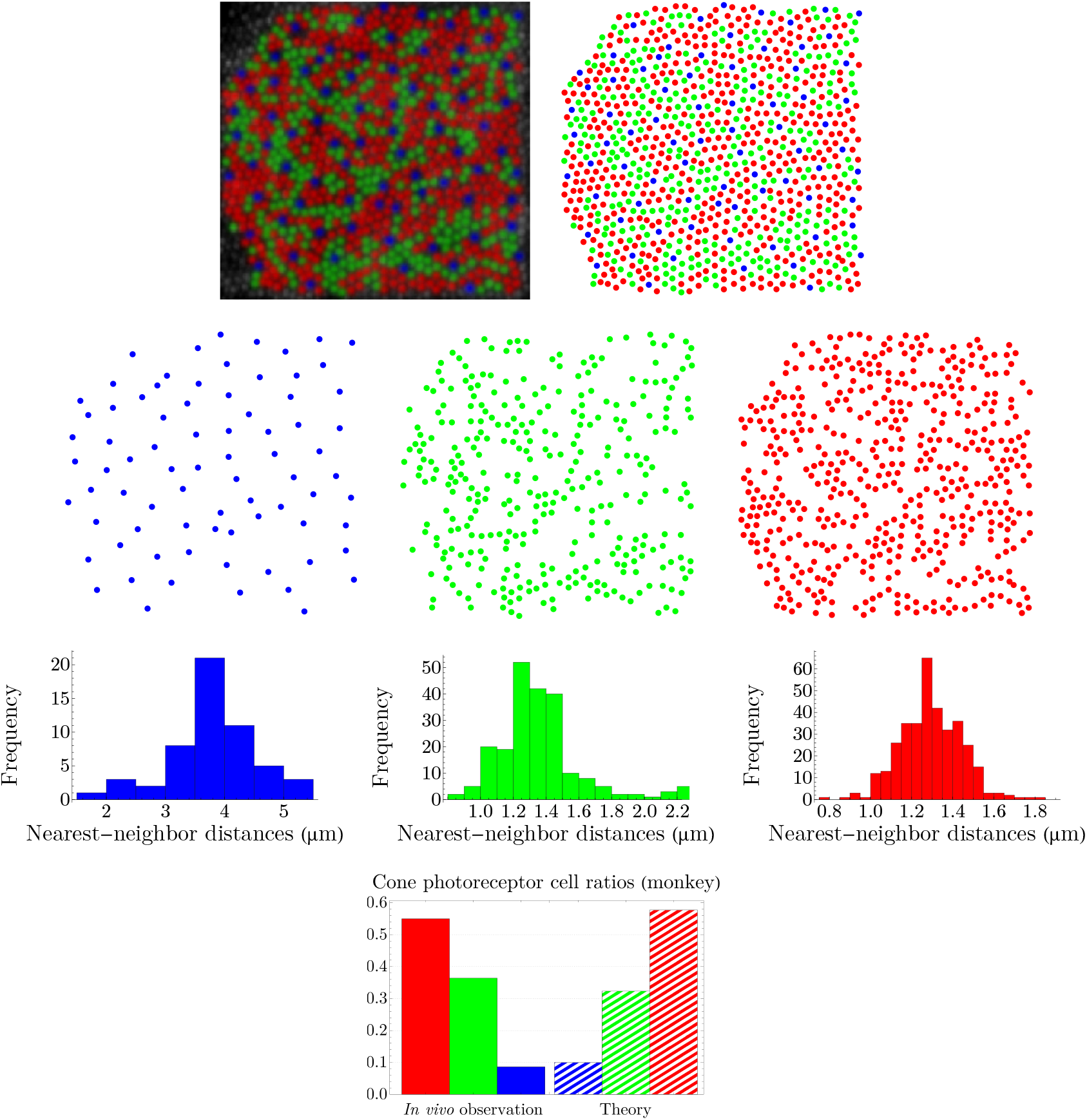
The image in the top-left corner, which shows the spatial distribution of cone photoreceptors in the nasal retina of a monkey (macaque), is provided by A. Roorda [36]. The entropies of the NND distributions in the third row read *h*_*b*_ = 1.019, *h*_*g*_ = 0.018, and *h*_*r*_ = −0.476. The predictions of the theory, illustrated in the fourth row, are evaluated at *β* = 1.174.

**Figure 8.**
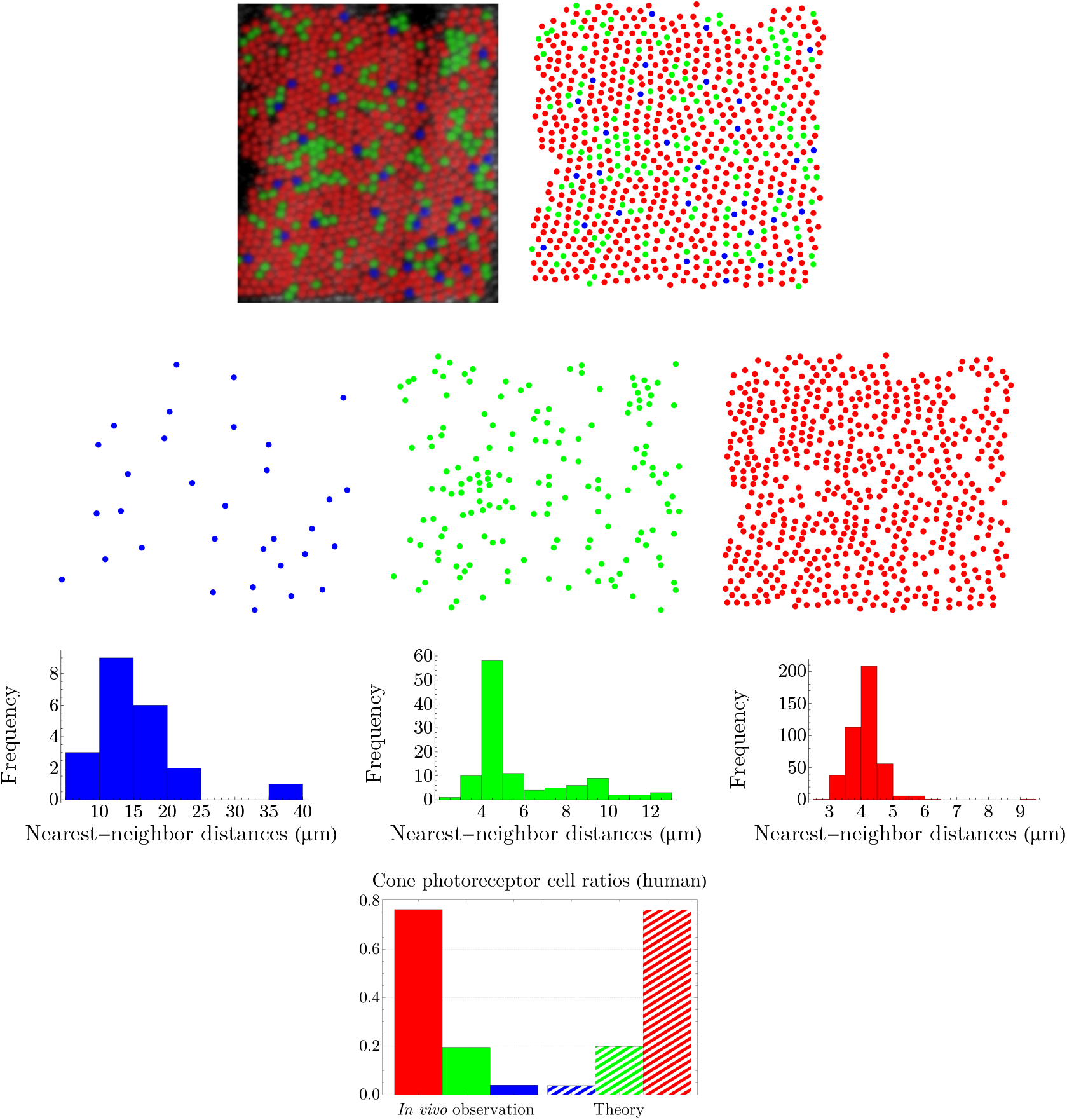
The image in the top-left corner, provided by A. Roorda [36], illustrates the spatial distribution of cone photoreceptors in the temporal retina of a human. The entropies of the NND distributions are: *h*_*b*_ = 2.977, *h*_*g*_ = 1.691, and *h*_*r*_ = 0.651; and *β* = 1.291.

**Figure 9.**
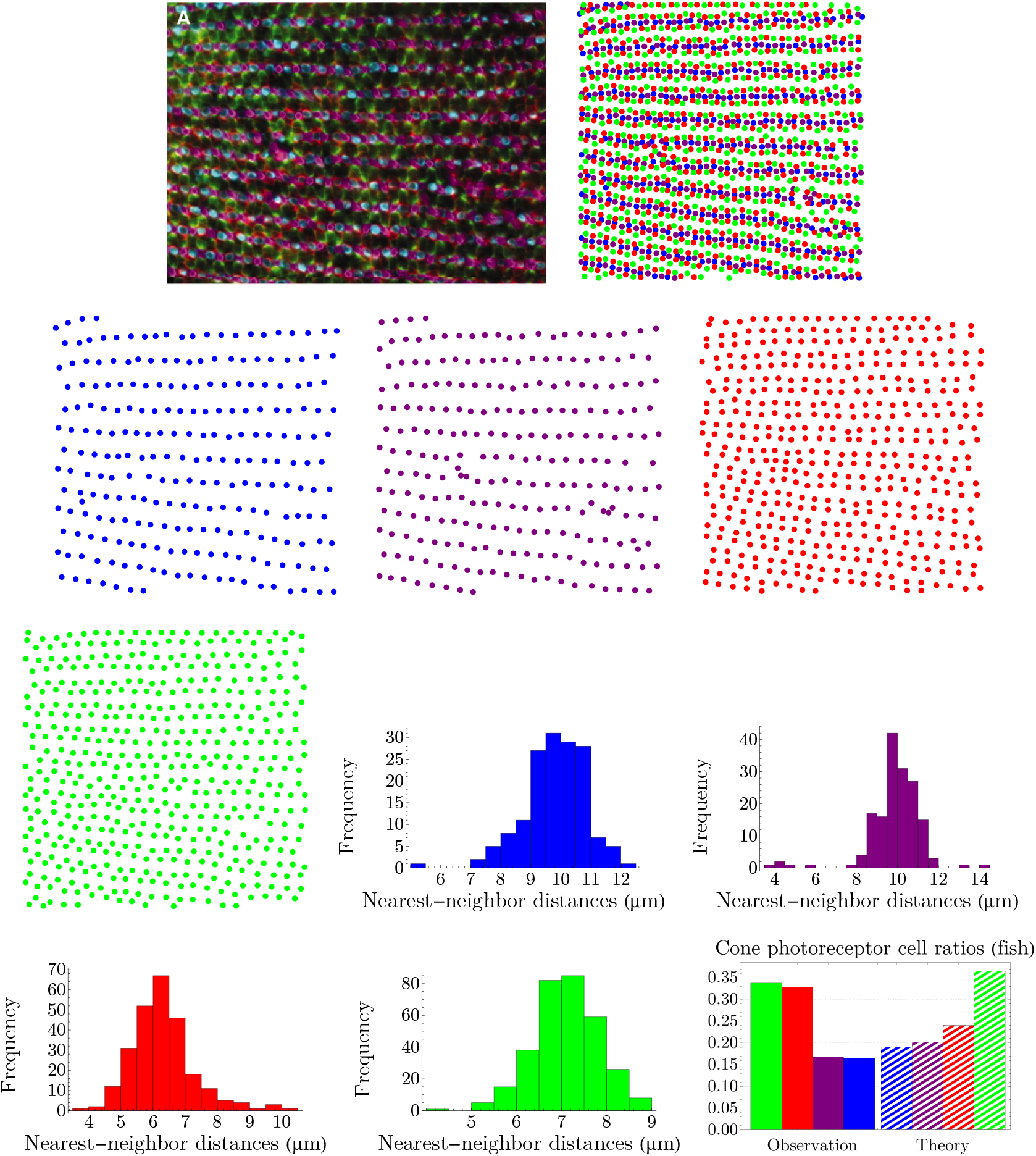
The image in the top-left corner, adapted from [37], shows the spatial distribution of cone photoreceptors in the retina of the zebrafish. The entropies of the NND distributions are: *h*_*b*_ = 1.471, *h*_UV_ = 1.440, *h*_*r*_ = 1.350, and *h*_*g*_ = 1.128; and *β* = 1.894.

**Figure 10.**
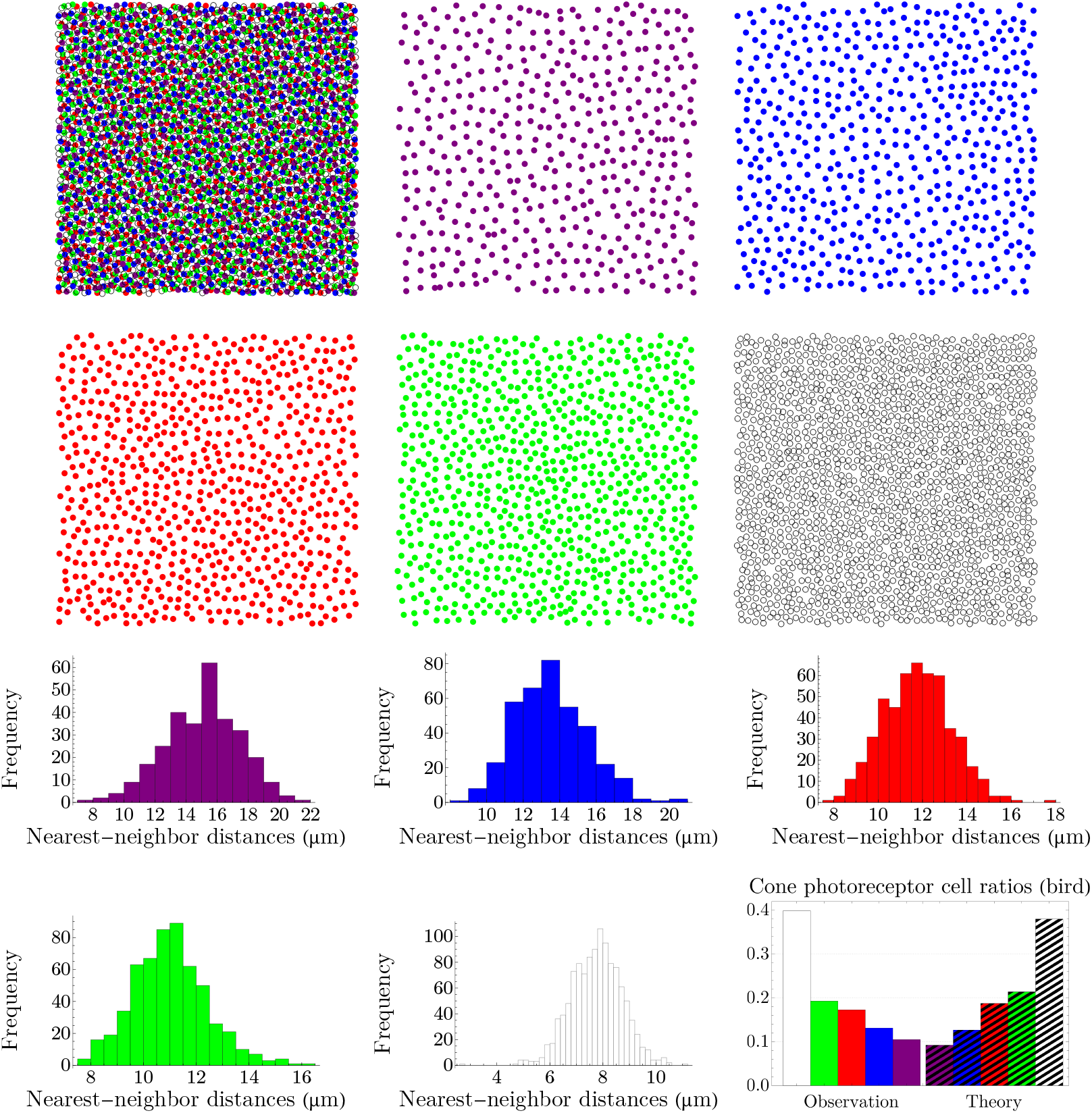
The digitized image of the spatial distribution of cone photoreceptors in the dorsal nasal retina of the chicken, shown in the top-left corner, is constructed from the data reported in [30]; double cones are represented as white. The entropies of the NND distributions read *h*_*v*_ = 2.291, *h*_*b*_ = 2.081, *h*_*r*_ = 1.826, *h*_*g*_ = 1.739, and *h*_*d*_ = 1.364; and *β* = 1.527.

### 3.3. Bounds on Retinal Coldness

We are in a position to address the issue raised at the end of Section 2: although we lack the knowledge of the numerical value 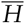 of *H*(*Y*| *S*); however, we have considered it as crucial information about the system and have represented it in terms of the Lagrange multiplier *β* (i.e., retinal coldness). In this subsection, we study the bounds on the value of 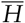, which lead to the estimation of *β*. To this end, we use the fact that for a given cone subtype, the distribution of the nearest-neighbor distances is peaked—see figures in Section 3.2—and thus, it can be approximated by a Gaussian.

We consider the nearest-neighbor distances as random variables, *X*_*s*_, where 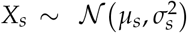 and *s* denotes cone subtypes. Note that in the case of a normal distribution, where the differential entropy is: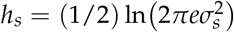 Equation (4) becomes:

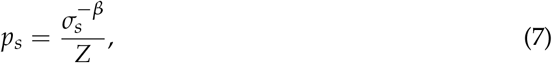

where 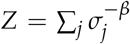. To estimate the bounds on the value of entropy, first, we define the random variable *W* as *W* = ∑_*s*_ *π*_*s*_*X*_*s*_, where *π*_*s*_ is the weight of the contribution of each cone subtype (i.e., the frequency of the appearance) and ∑_*s*_ *π*_*s*_ = 1. Since *W* has a normal distribution, its entropy is related to its variance, *σ*^2^, as (1/2) ln *σ*^2^ + const. Thus, we can obtain the lower and upper bounds of the entropy by minimizing and maximizing the variance, respectively. In the following, we study these two extreme cases.

Variance of *W* reads 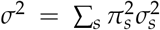. By the method of Lagrange multipliers, we minimize *σ*^2^, subjected to the constraint ∑_*s*_ *π*_*s*_ = 1. It turns out that [38], for 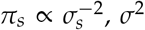 is minimized, which implies the minimization of the entropy of *W*. By comparing this *π*_*s*_ with (7), we establish the upper bound of *β* as 2. To obtain the lower bound of *β*, we maximize the entropy of *W*, which implies the maximization of *σ*^2^, subjected to ∑_*s*_ *π*_*s*_ = 1. This happens by letting *π*_*s*_ corresponding to the largest *σ*_*s*_ to be 1 and all the other *π*_*s*_’s vanish. This scenario is not desirable, as contributions of various colors vanish. To obtain an acceptable maximum value for the entropy of *W*, we consider the uncertainties associated with random variables *π*_*s*_*X*_*s*_ to be equal, that is, we equalize variances of *π*_*s*_*X*_*s*_ by Considering 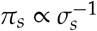, which results in *σ*^2^ = *C*^2^ ∑_*s*_ = *C*^2^*N. C* is a proportionality constant, i.e., 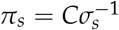, and *N* is the number of cone subtypes. By comparing this *π*_*s*_ with (7), we establish the lower bound of *β* as 1.

Among species studied in Section 3.2, fish and bird have more ordered retinal cone mosaics, where in the former is highly regular and in the latter is semi-random [16]. These two species’ corresponding *β*’s are more closer to 2 than the other species. Thus, more ordered patterns correspond to a lower retinal temperature or a higher coldness, as expected in thermodynamics. More irregular mosaics—like in rodent, dog, monkey, or human—have higher retinal temperatures. Overall, for vertebrate retinas, under the assumption that the NND distributions are peaked, we always have: 1 < *β* < 2.

## 4. Lemaître’s Law

Lemaître’s law is the virial equation of state for two-dimensional cellular networks, which relates two measures of disorder (i.e., thermodynamic variables), namely, the fraction of hexagons to the width of the polygon distribution [11,19,20,39–41]. Although at first proposed for two-dimensional foams, it has been shown that a wide range of planar cellular networks in nature obeys Lemaître’s law, ranging from biology such as avian cones [30], epithelial cells [42], and mammalian corneal endothelium [43], to physics such as amorphous graphene [41], the Ising model [44], Bénard–Marangoni convection [45], silicon nanofoams [46], and silica bilayers [47]. It can be obtained by maximizing the entropy, *H* = − ∑_*n*_ ≥_3_ *p*_*n*_ ln *p*_*n*_, where *p*_*n*_ is the probability, or the frequency of the appearance, of an *n*-sided polygon, while considering the following information:

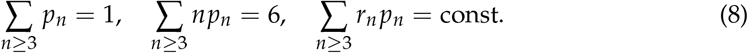

The first relation is the normalization condition, and the second one is a consequence of Euler’s relation concerning the topology of the structure, which assumes only three lines meet at a vertex. Networks with higher vertices can be transformed into trivalent vertices by appropriate transformations [48]. The function *r*_*n*_ in the last relation depends on the geometry or the underlying dynamics of cells (polygons). Lemaître and colleagues assumed *r*_*n*_ = 1/*n* as an empirical observation made by measuring the areas of cells in a two-dimensional mosaic produced by hard discs moving on an air table [19,20]. At first glance, the choice of *r*_*n*_ = 1/*n* seems not applicable in a general setting. Indeed, it was already mentioned in [20] that this particular form of *r*_*n*_ cannot be valid for all cellular mosaics, as, for instance, it is incompatible with the well-known Lewis’ law [49], which assumes that the average area of polygons is linear in *n*. However, the authors of [20] speculated that the remarkable universality of Lemaître’s law suggests that the particular choice of *r*_*n*_ = 1/*n* has probably a deeper meaning than expected.

Without considering any ad hoc constraint, we derive Lemaître’s law as a special case of our formalism explained in Section 2. To this end, first, we generalize the Lagrangian introduced in Equation (3) as [13],

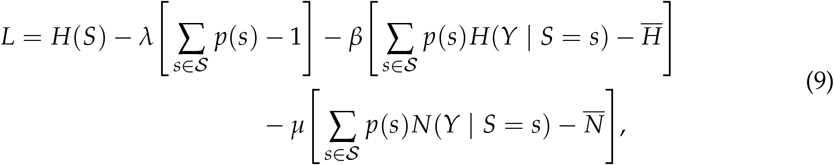

where we have assumed the following additional information: 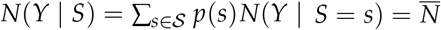, which *N*(*Y* |*S*) is the average number of cells in local environment and *µ* is a Lagrange multiplier. By solving *∂L*/*∂p*(*s*) = 0, we obtain:

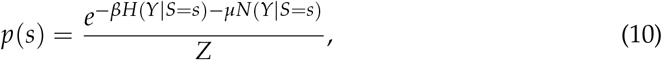

where *Z* = ∑_*w∈*_ *𝒮* exp [−*βH*(*Y* | *S* = *w*) − *µN*(*Y*| *S* = *w*)]. We simplify the notations in (10) and write:

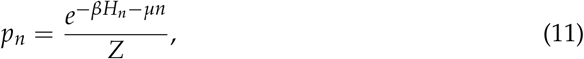

where we have replaced *s* by *n. p*_*n*_ is the probability of having an *n*-sided polygon, or its frequency of appearance, and *Z* = ∑_*n≥* 3_ exp[−*βH*_*n*_ −*µn*]. To calculate *H*_*n*_, we consider a general standardized discrete distribution, which its density can be expanded as [50],

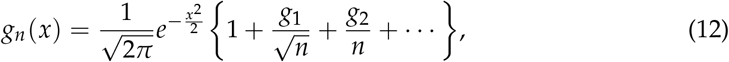

with 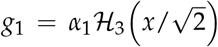 and 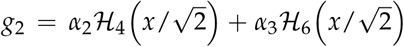, where *α*_1_, *α*_2_, *α*_3_ are constants and ℋ _*k*_(·) is the *k*th Hermite polynomial. Note that as *n* → ∞, *g*_*n*_(*x*) approaches the standard normal distribution. Now that we have *g*_*n*_(*x*) at our disposal, we can calculate, its differential entropy, 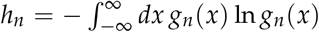 ln *g*_*n*_(*x*). Since ℋ_3_(*x*) = 8*x*^3^ − 12*x* is an odd function of *x* its integral vanishes, and thus, the first nonzero correction term is of the order 1/*n*. We obtain:

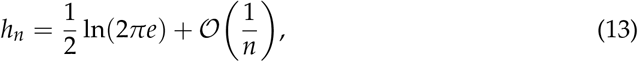

where the first term is the entropy of the standard normal distribution. By plugging (13) into (11), we arrive at

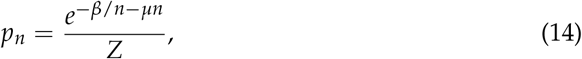

where *Z* = ∑_*n*_ ≥_3_ exp [−*β*/*n* − *µn*] and we have absorbed the constants included in 𝒪 (1/*n*) in *β*. Equation (14) sheds light on the origin of *r*_*n*_ = 1/*n*, which Lemaître and colleagues had obtained for a specific two-dimensional mosaic [19,20]. Since the calculations leading to (14) only assume a general discrete distribution, the universality of Lemaître’s law becomes evident.

The variance, *µ*_2_, of the distribution *p*_*n*_ in (14) reads

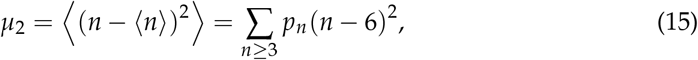

where we have used Euler’s relation, ⟨*n*⟩ = 6. The second moment of *p*_*n*_, *µ*_2_, demonstrates a deviation from the hexagonal configuration and can be interpreted as a measure of topological disorder. By exploiting (14) and (15), Lemaître’s law, as a relation between two measures of disorder, *µ*_2_ and *p*_6_, has been obtained as [11,19,20,41,46],

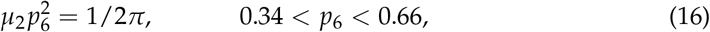

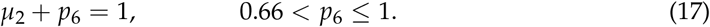

We present a simple and intuitive derivation of (16) and (17), which is inspired and developed in discussion with C. Beenakker and I. Pinelis [51]. For (16) to hold, *p*_6_ should be large and thus *p*_*n*_ in (14) should peak at *n* = 6. This allows us to approximate *p*_*n*_ near *n* = 6 by a normal distribution, *P*_*n*_, centered at *n* = 6 while ignoring the discreteness of *n*. We can let *n* vary from −∞ to ∞ since only those *n*’s close to the peak have notable contributions, provided that *p*_6_ is not too small. Thus, we have: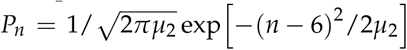,which results in 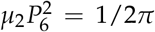. For (17) to hold, the probabilities *p*_*n*_’s for *n* ∉{ 5, 6, 7} should be negligible compared to *p*_*n*_’s for *n*∈ {5, 6, 7 }; as a result, the discreteness of *n* cannot be neglected in this case, since only three *n*’s contribute. The constraint *n* = 6 implies that *p*_*n*_ should *sharply* peak at *n* = 6, leading to *µ*_2_ →0 as *p*_6_ →1, and thus: *µ*_2_ + *p*_6_ → 1. Note that although in (9), we have assumed information about seemingly unrelated quantities *H*(*Y* |*S*) and *N*(*Y* | *S*) represented in terms of their corresponding Lagrange multipliers *β* and *µ*, the peakedness of *p*_*n*_ and thus *h*(*n*) *β*/*n µn* at *n* = 6 gives us a relation between *β* and *µ*. Since *h*^′^(6) = *β*/6^2^ − *µ* = 0, we have: *β* = 36*µ*.

To obtain regions of validity of 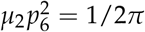 and *µ*_2_ + *p*_6_ = 1, numerical analyses are performed and the results are shown in Figure 11. The left panel illustrates *µ*_2_ as a function of *p*_6_, where red points are obtained from (14), subjected to the constraint ⟨*n*⟩ = 6, and the dashed, blue and yellow curves correspond to 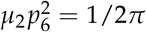 and *µ*_2_ + *p*_6_ = 1, respectively. Simulations suggest that the known lower bound of (16) can be relaxed to 0.27, that is,

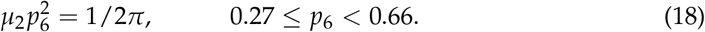

**Figure 11.**
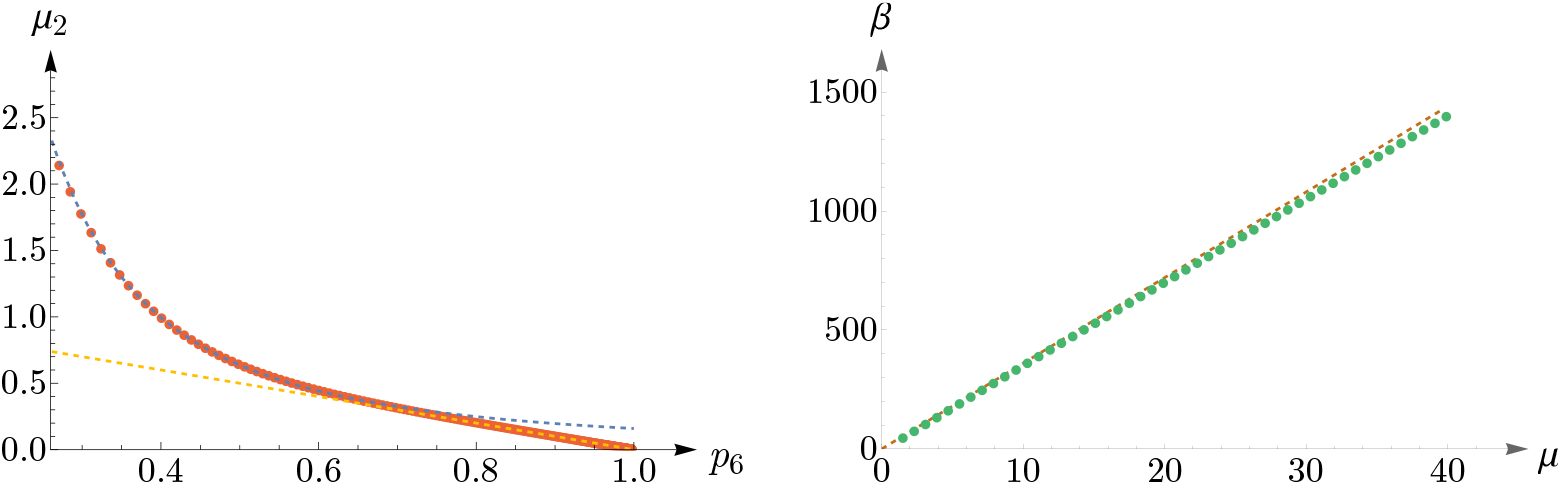
In the left panel, the dashed, blue and yellow curves correspond to 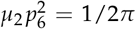 and *µ*_2_ + *p*_6_ = 1, respectively. Red points are obtained from (14), subjected to the constraint ⟨*n*⟩ = 6. This plot suggests that the known lower bound of (16) can be relaxed to 0.27. The right panel presents a comparison between the analytical result of *β* = 36*µ*, shown as a dashed brown curve, and the values of (*µ, β*), shown as green points, obtained from (14).

In the right panel of Figure 11, we have shown *β* as a function of *µ*, where the dashed, brown curve represents *β* = 36*µ* and the green points depict the values of (*µ, β*) obtained from (14), subjected to the constraint ⟨*n*⟩ = 6.

As *p*_6_ decreases, going from 0.27 to 0.25, the peak of *p*_*n*_ shifts from *n* = 6 to *n* = 5 and remains so up to *p*_6_ = 0.16, see the left panel of Figure 12. Again, we can approximate *p*_*n*_ by a Gaussian, which this time peaks at *n* = 5. In the right panel of Figure 12, we have shown the values of (*p*_5_, *µ*_2_), obtained from (14) and subjected to the constraint ⟨*n*⟩ = 6, in red points, and the Gaussian as a dashed, blue curve.

**Figure 12.**
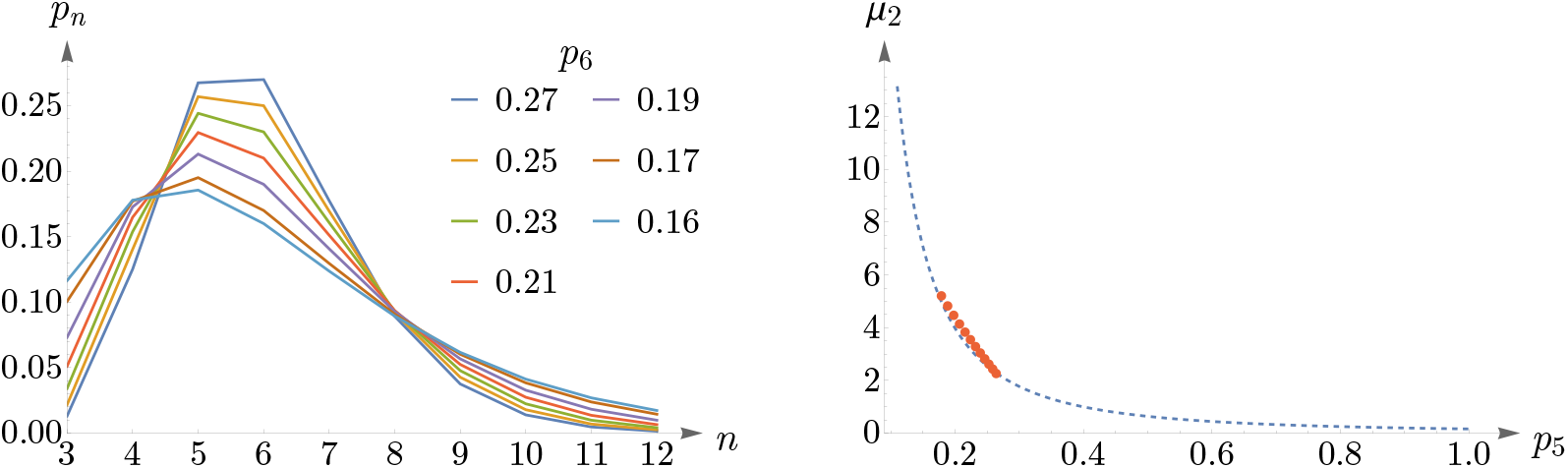
The left panel shows the mode shift of *p*_*n*_ in (14) from *n* = 6 to *n* = 5 as *p*_6_ decreases from 0.27 to 0.25. In the right panel, the red points depict (*p*_5_, *µ*_2_) obtained from *p*_*n*_ and subjected to the constraint ⟨*n*⟩ = 6, and the Gaussian is shown as a dashed, blue curve.

By decreasing *p*_6_ further, the peak shifts from *n* = 5 to *n* = 4, and eventually, *p*_*n*_ becomes monotonically decreasing, see the left panel of Figure 13. For small values of *p*_6_, going from 0.09 to 0.07, *p*_*n*_ becomes a U-shaped distribution, as is shown in the right panel of Figure 13.

**Figure 13.**
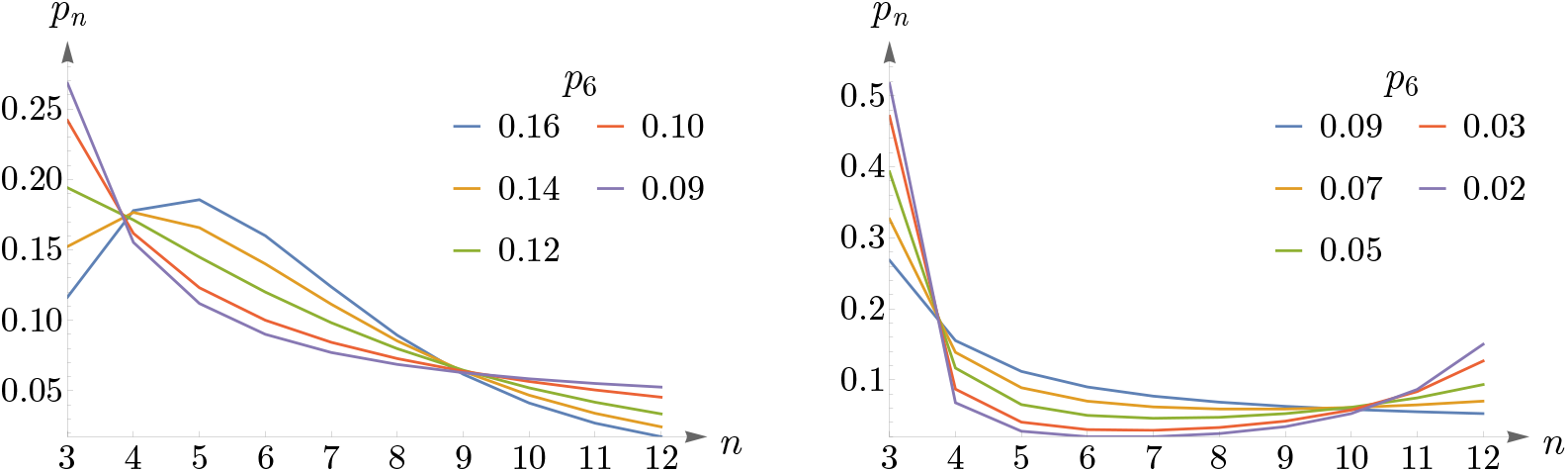
Left panel shows that as *p*_6_ decreases, the peak of *p*_*n*_ shifts from *n* = 5 to *n* = 4, and eventually, *p*_*n*_ becomes a monotonically decreasing distribution. The right panel depicts a change in the shape of *p*_*n*_ from a monotonically decreasing to a U-shaped distribution for small values of *p*_6_. In both panels, it is assumed that ⟨*n*⟩ = 6.

Most two-dimensional cellular networks in nature have an abundance of hexagons, and they likely obey (17) and (18). Low values of *p*_6_ may correspond to amorphous or artificially generated networks. In the following, we examine several cases of mosaics that are artificially generated: random fragmentation, Feynman diagrams, the Poisson network, and semi-regular Archimedean tiling. We demonstrate that all these networks still obey *p*_*n*_ in (14).

In [52], specific artificial, two-dimensional cellular structures are generated by a fragmentation process. One way to construct these networks is by a random selection of a cell among all cells, and then this cell is to be fragmented into two cells by adding an edge randomly. The side number distribution of cells in this system is obtained by a mean-field model as [52],

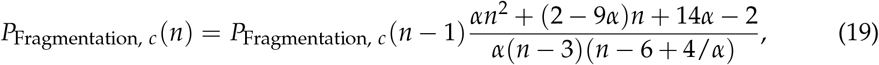

where *α* = 0.356 and *P*_Fragmentation, *c*_(6) = 0.125. Equation (19) can be solved as

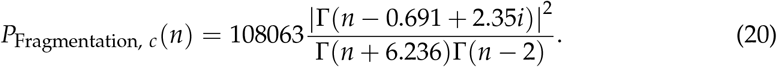

In the top-left corner of Figure 14, we have shown *P*_Fragmentation, *c*_(*n*) as a blue curve and *p*_*n*_ in (14) as a dashed, orange curve.

**Figure 14.**
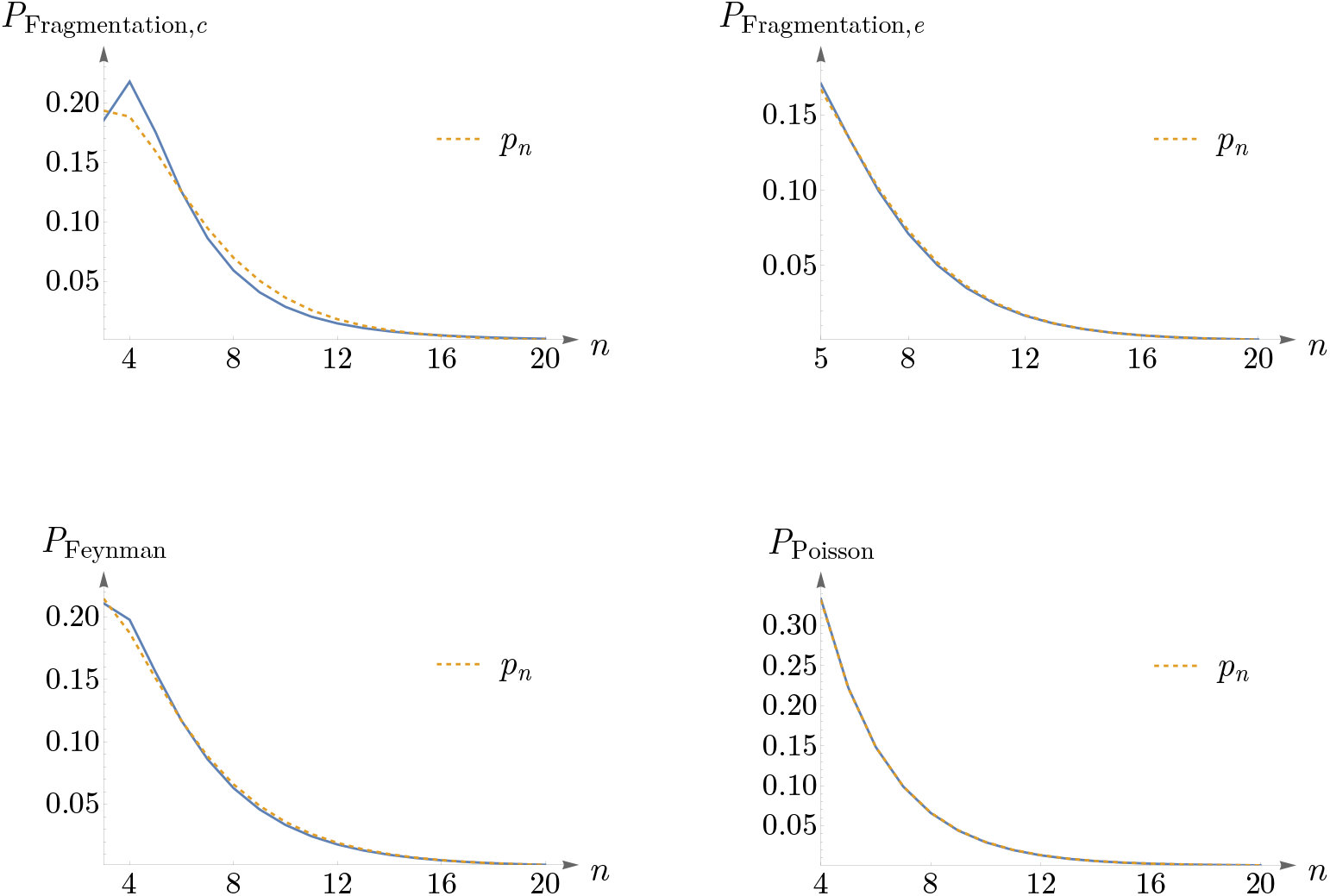
Four artificially generated networks, by random fragmentation, Feynman diagrams, and the Poisson network, are compared to the probability distribution *p*_*n*_ in (14) with the constraint ⟨*n*⟩ = 6.

Another way to construct such networks is by a random selection of an edge among all cell edges followed by selecting one of the cells which shares this edge, and then this cell is to be fragmented into two cells as in the previous case [52]. The probability distribution of the number of cell sides reads [52],

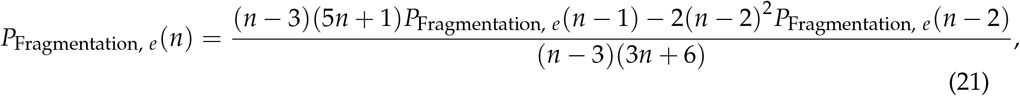

with *P*_Fragmentation, *e*_(4) = 0.196 and *P*_Fragmentation, *e*_(6) = 0.134. In the top-right corner of Figure 14, a comparison between *P*_Fragmentation, *e*_(*n*) and *p*_*n*_ is shown.

The ensemble of planar Feynman diagrams with a cubic interaction (i.e., planar *ϕ*^3^ diagrams with a fixed number of vertices) is equivalent to the ensemble of polygons with trivalent vertices [53,54]. The probability distribution of the number of cell edges is obtained as [53,54],

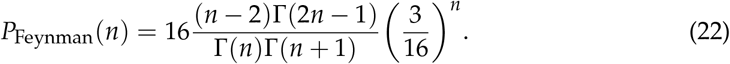

See the bottom-left corner of Figure 14, for a comparison between *P*_Feynman_(*n*) and *p*_*n*_.

The two-dimensional Poisson network studied in [55] can be obtained from a tessellation of a surface based on Poisson point distribution. The distribution of the number of cell sides reads [55],

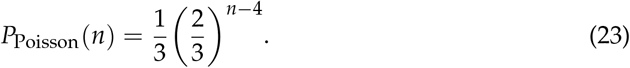

A comparison between *P*_Poisson_(*n*) and *p*_*n*_ is shown in the bottom-right corner of Figure 14.

The Archimedean tilings, obtained by Kepler, are the analogs of the Archimedean solids. Eight of them are semi-regular and consist of regular polygons at each vertex [56]. In the left panel of Figure 15, we have shown one of these semi-regular tilings, known as truncated hexagonal tiling, consisting of two dodecagons and one triangle at each vertex. The right panel of Figure 15 shows *p*_*n*_ in (14) as *p*_6_ → 0 and *n* = ⟨6⟩. This plot corresponds to a pattern that comprises an abundance of triangles with dodecagons amongst them and is in agreement with truncated hexagonal tiling.

**Figure 15.**
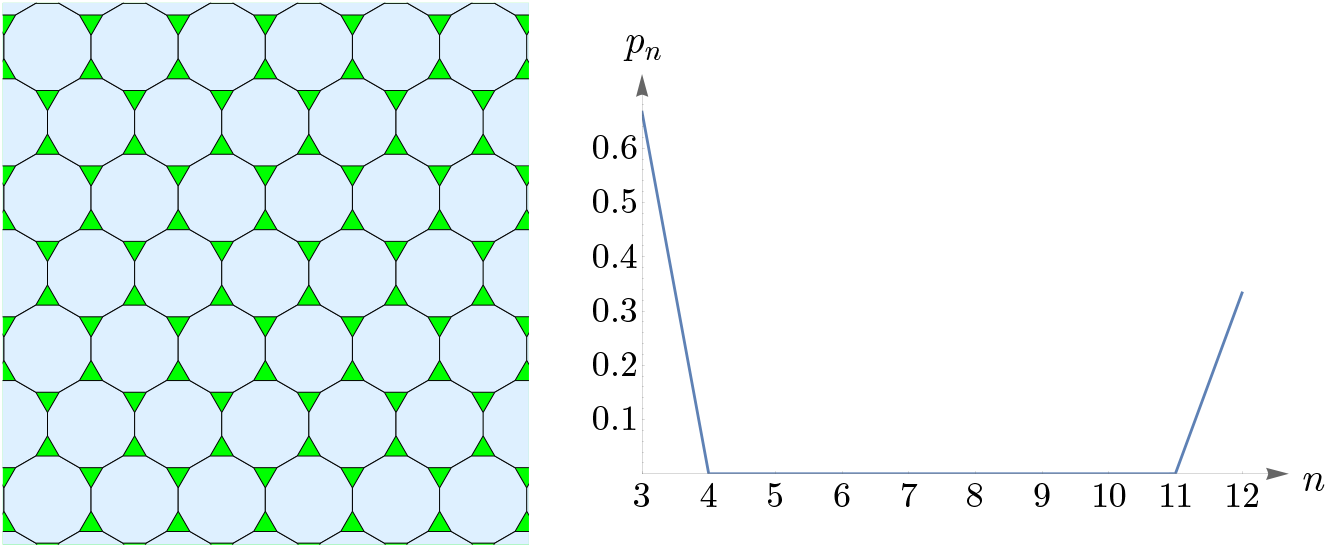
Left panel shows truncated hexagonal tiling, with ≈ 66% triangles and ≈ 34% dodecagons. The right panel shows *p*_*n*_ in (14) as *p*_6_ → 0 and ⟨*n*⟩ = 6.

### 4.1. Human Cone Mosaics

In this subsection, we examine Lemaître’s law in the case of the human retina, which can be viewed as a natural, two-dimensional cellular network. To partition the retinal field of Figure 1 into polygons, we construct the corresponding Voronoi tessellation. Each Voronoi polygon is generated by a cone cell in a way all points in a given polygon are closer to its creating cone cell than to any other [57]. In the top row of Figure 16, we have shown Voronoi tessellations of the spatial arrangements of blue, green, and red cones in a living human retina. At the bottom, the Voronoi tessellation of the whole pattern of cones is presented. The fractions of *n*-sided bounded polygons are reported in the figure caption.

**Figure 16.**
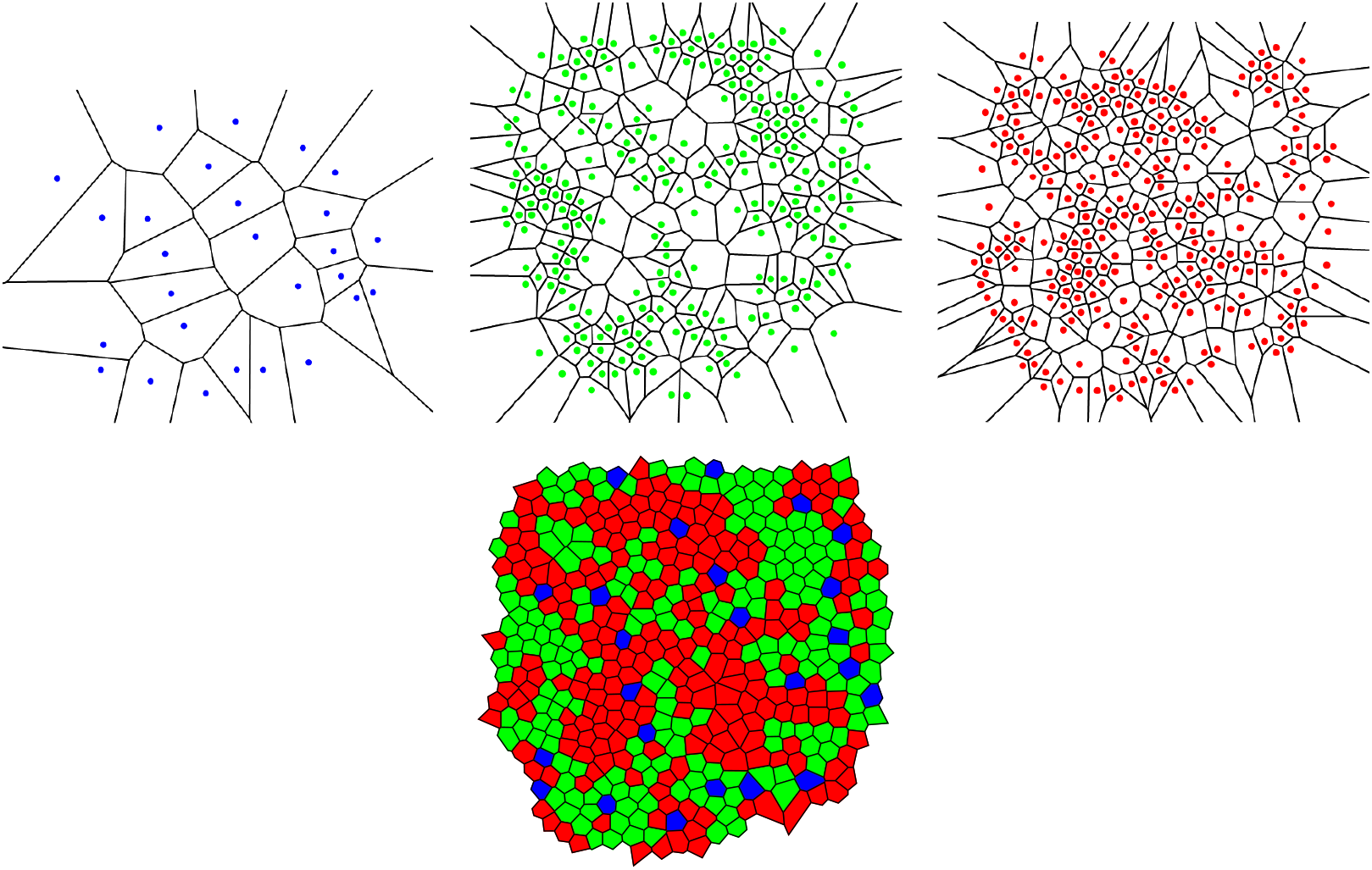
In the top row, we have shown Voronoi tessellations of the three cone photoreceptor subtypes in a living human retina in Figure 1. The fraction of *n*-sided bounded polygons, 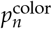 in each case reads 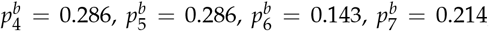 and 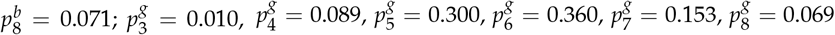 and 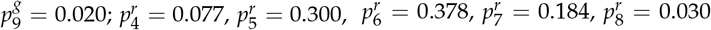 and 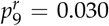.The Voronoi tessellation of the whole retinal field is illustrated at the bottom, with the fractions of *n*-sided polygons as *p*_4_ = 0.012, *p*_5_ = 0.171, *p*_6_ = 0.718, *p*_7_ = 0.086, and *p*_8_ = 0.012.

If we assume a high value of *p*_6_ indicates the regularity of the corresponding cone mosaic, Figure 16 demonstrates that the spatial arrangement of blue cones is more random than those of green and red cones, where for blue cones we have: 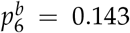 while 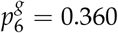 and 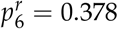 for greens and reds, respectively. This finding is in agreement with [28]. Note that, as is shown at the bottom of Figure 16, in contrast to the cone subtypes, the whole spatial arrangement of human cones is highly ordered, with *p*_6_ = 0.718.

We have shown Lemaître’s law as applied to human cone mosaics in Figure 17. In the left panel—the case of blue cone mosaic—the experimental value of 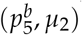 is depicted as a blue point, and the dashed, dark-gray curve corresponds to 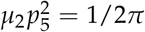 and the dashed, light-gray curve to *µ*_2_ + *p*_5_ = 1. The cases of greens, reds, and the entire pattern of cones (in black) are shown in the right panel.

**Figure 17.**
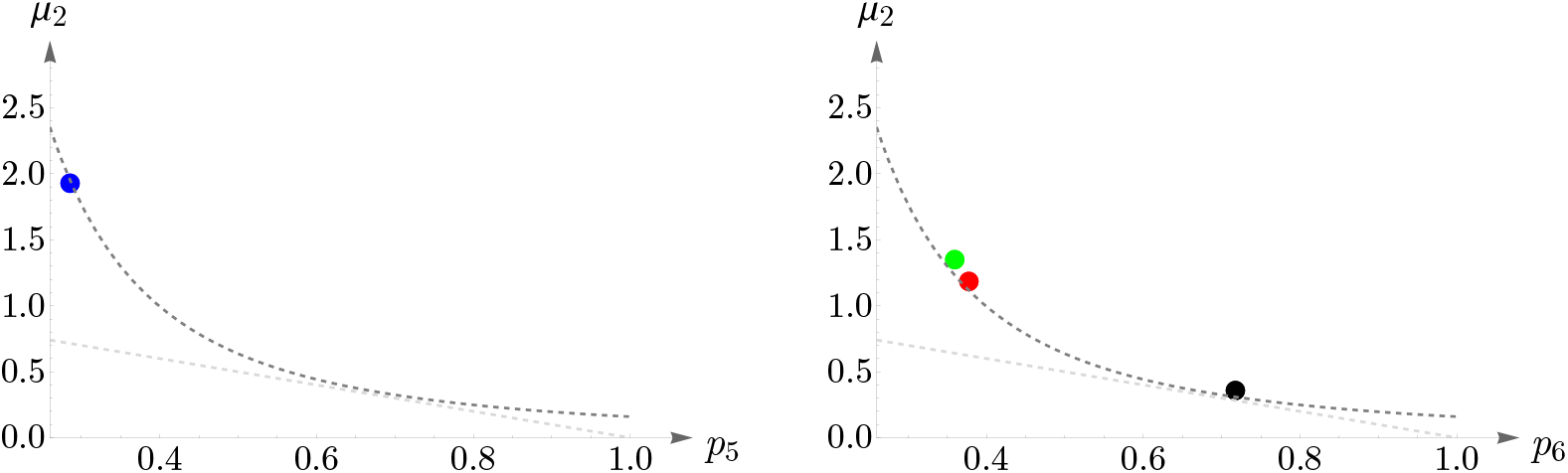
Blue, green, red, and black points depict the experimental values of 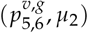, *n* = 5, 6, for human cone mosaics (the black point represents the whole pattern of cones). Lemaître’s law is shown as dashed, gray curves.

As another illustration, the behavior of cones in a different subject is shown in Figure 18. The image in the left panel, adapted from [58], shows human cone mosaics at six different retinal locations: two, four, six, eight, ten, and twelve degrees of retinal eccentricities, temporal to the fovea. The right panel shows the agreement between human cone mosaics and Lemaître’s law.

**Figure 18.**
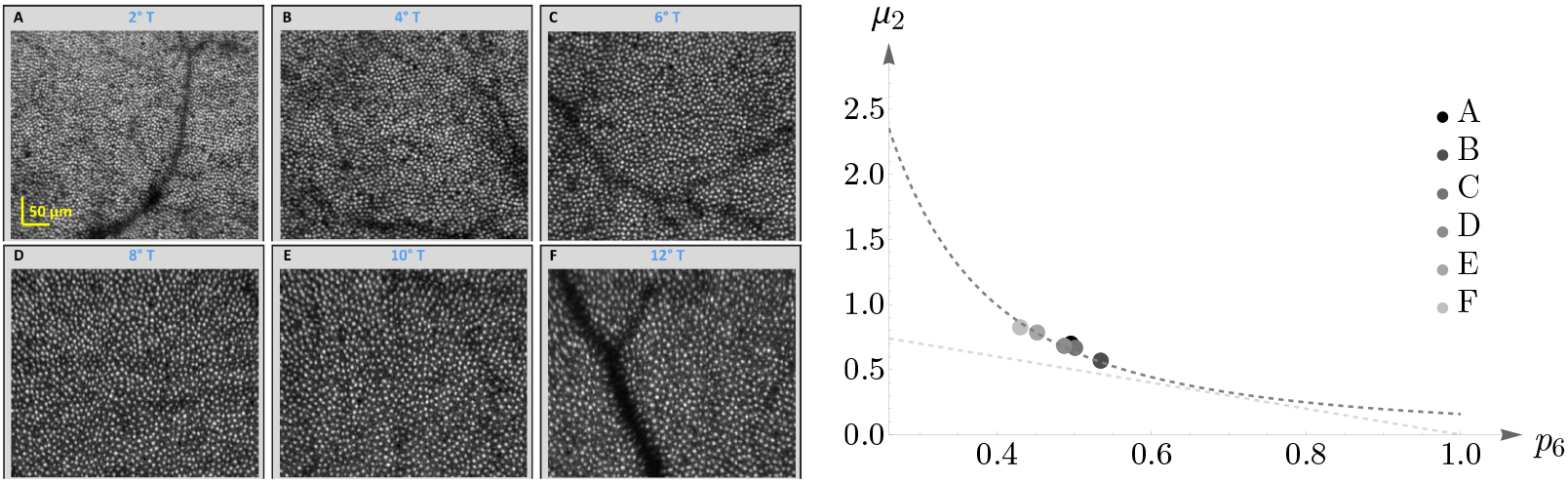
The left panel, adapted from [58], shows the spatial distributions of cone photoreceptors in the retina of a living human eye at a range of retinal eccentricities. In the right panel, we have depicted cones’ behavior—the whole pattern in each case—concerning Lemaître’s law.

### 4.2. Vertebrate Cone Mosaics: From Rodent to Bird

Here, we apply the approach of Section 4.1 to rodent, dog, monkey, human, fish, and bird. The results are summarized in Figures 19, 20, 21, 22, 23, and 24. In each case, the experimental value of 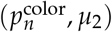 is depicted in the color of its respective cone subtype, and the black point represents the whole pattern of cones in a given retinal field.

**Figure 19.**
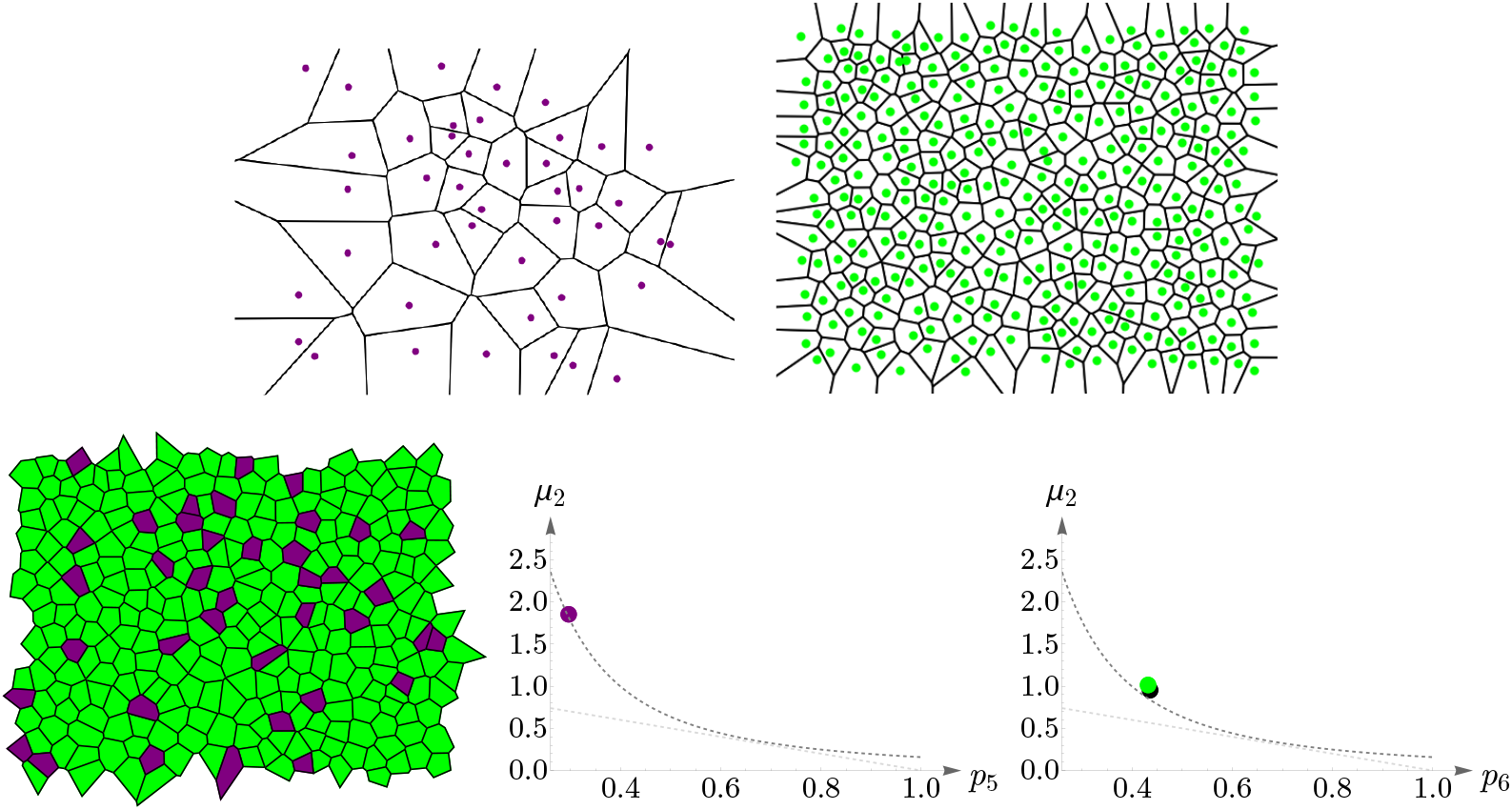
Voronoi tessellations of the rodent retinal field in Figure 5 and the corresponding cones’ behavior concerning Lemaître’s law. The violet and green points in the plots correspond to the experimental values of 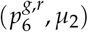 for short- and long-wavelength-sensitive cones, respectively. The black point corresponds to the whole retinal mosaic. Lemaître’s law is shown as dashed, gray curves.

**Figure 20.**
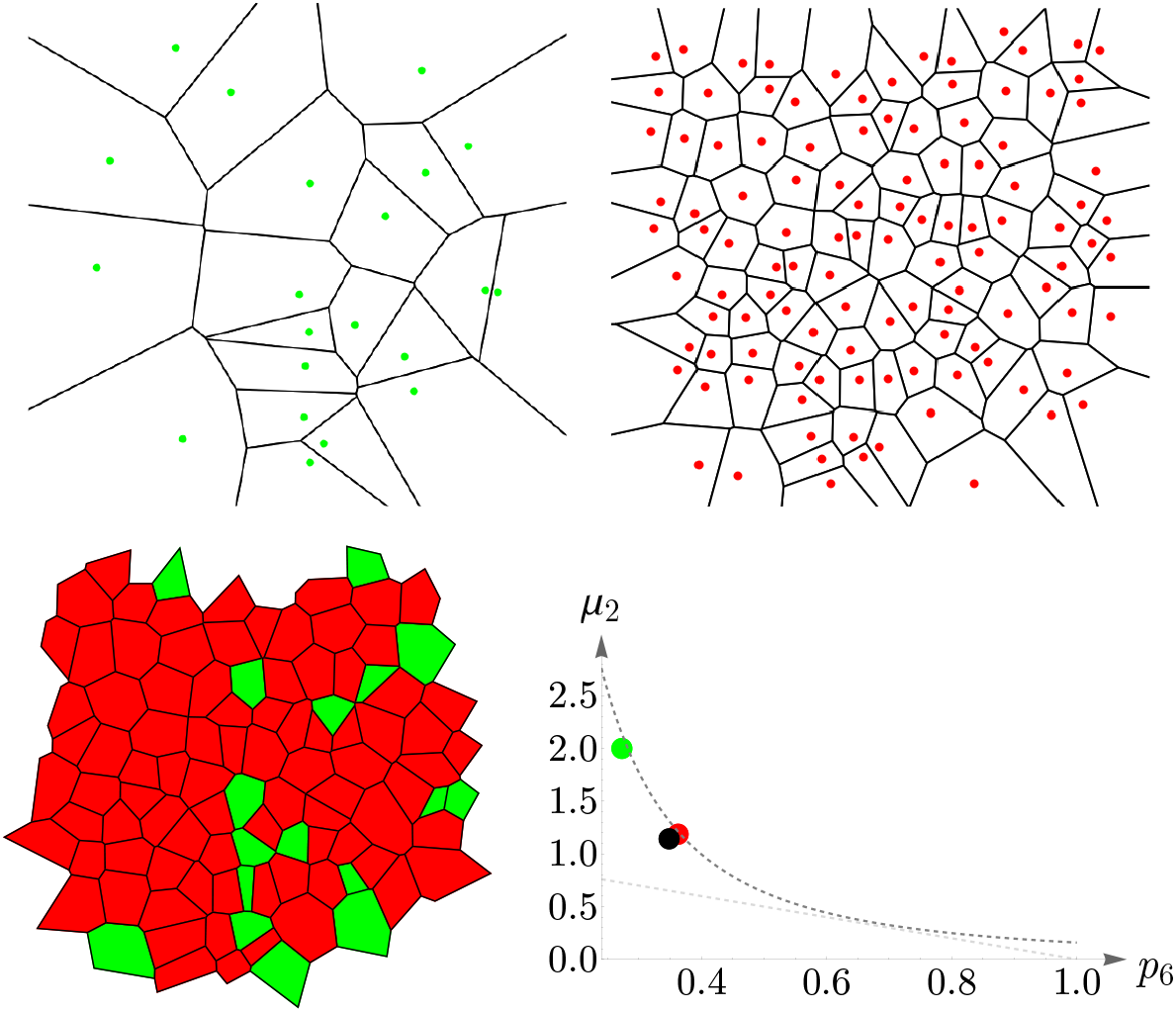
Voronoi tessellations of the dog retinal field in Figure 6 and the corresponding cones’ behavior concerning Lemaître’s law. The green and red points in the plot correspond to the experimental values of 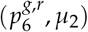 for short- and long-/medium-wavelength-sensitive cones, respectively. The black point corresponds to the whole retinal mosaic.

**Figure 21.**
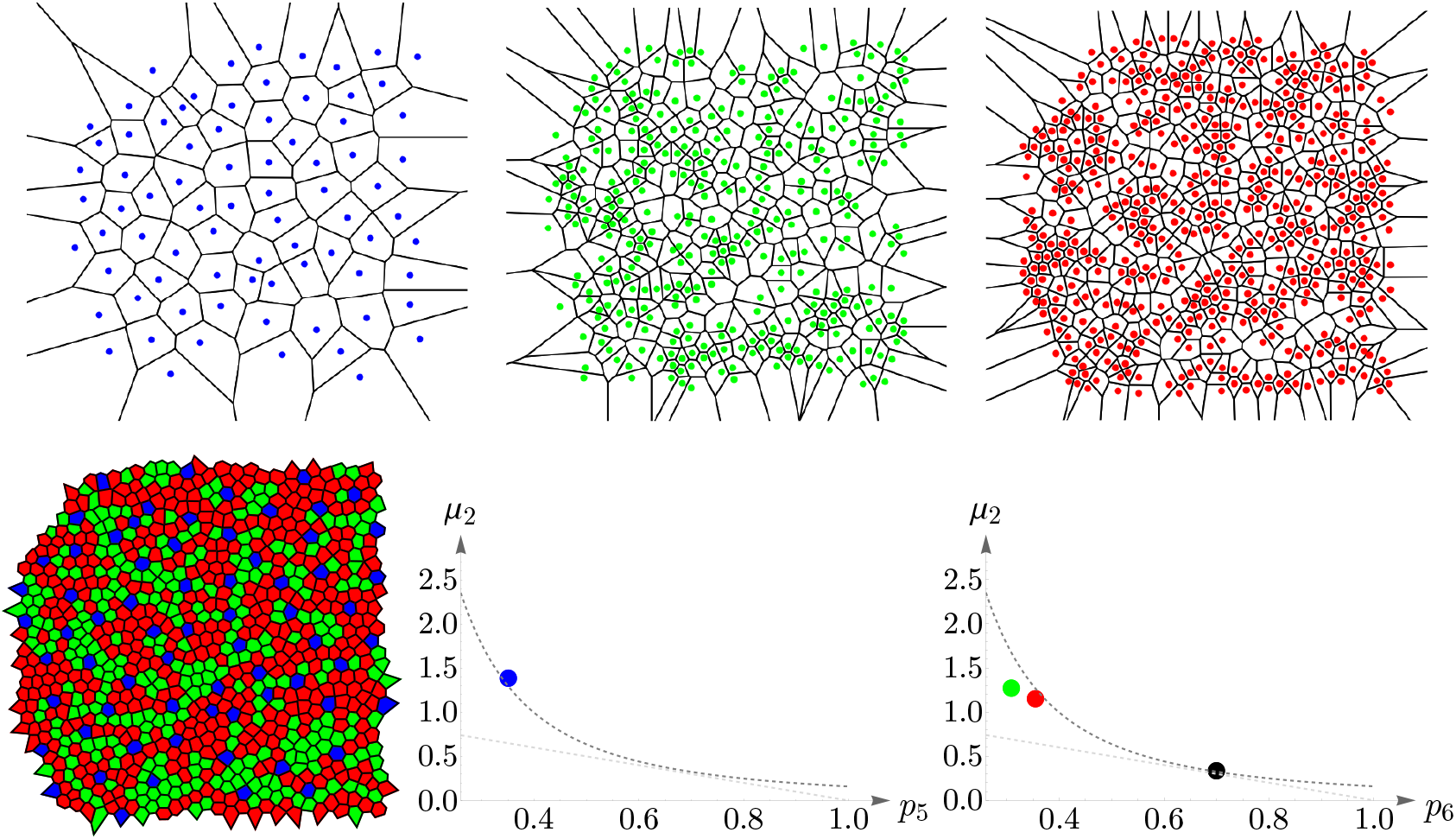
Voronoi tessellations of the monkey retinal field in Figure 7 and the corresponding cones’ behavior concerning Lemaître’s law. In the plots, points’ colors reflect the corresponding cone subtypes and the black point represents the whole retinal field.

**Figure 22.**
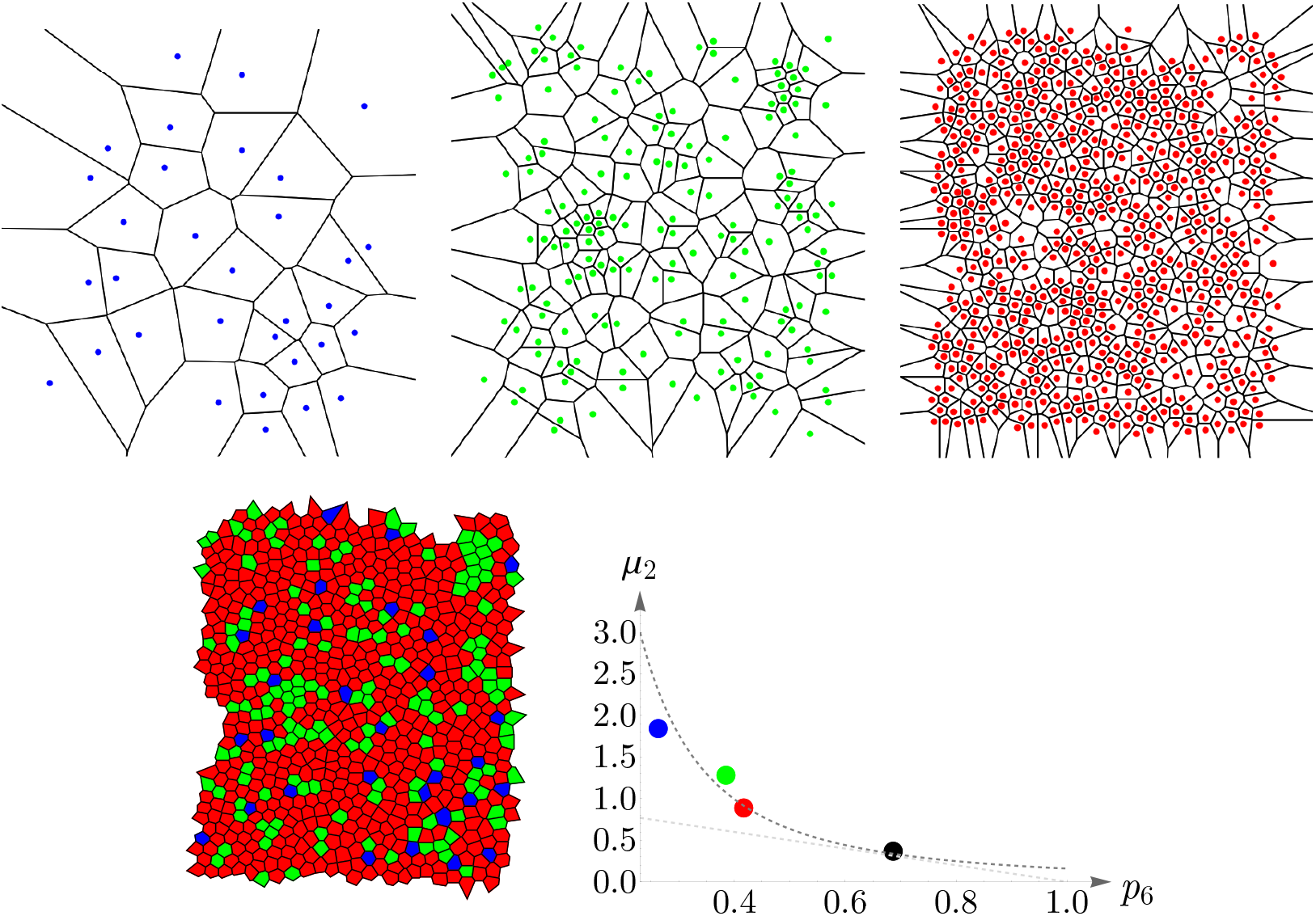
Voronoi tessellations of the human retinal field in Figure 8 and the corresponding cones’ behavior concerning Lemaître’s law. Points in colors in the plot correspond to the cone subtypes, and the black point represents the whole retinal field.

**Figure 23.**
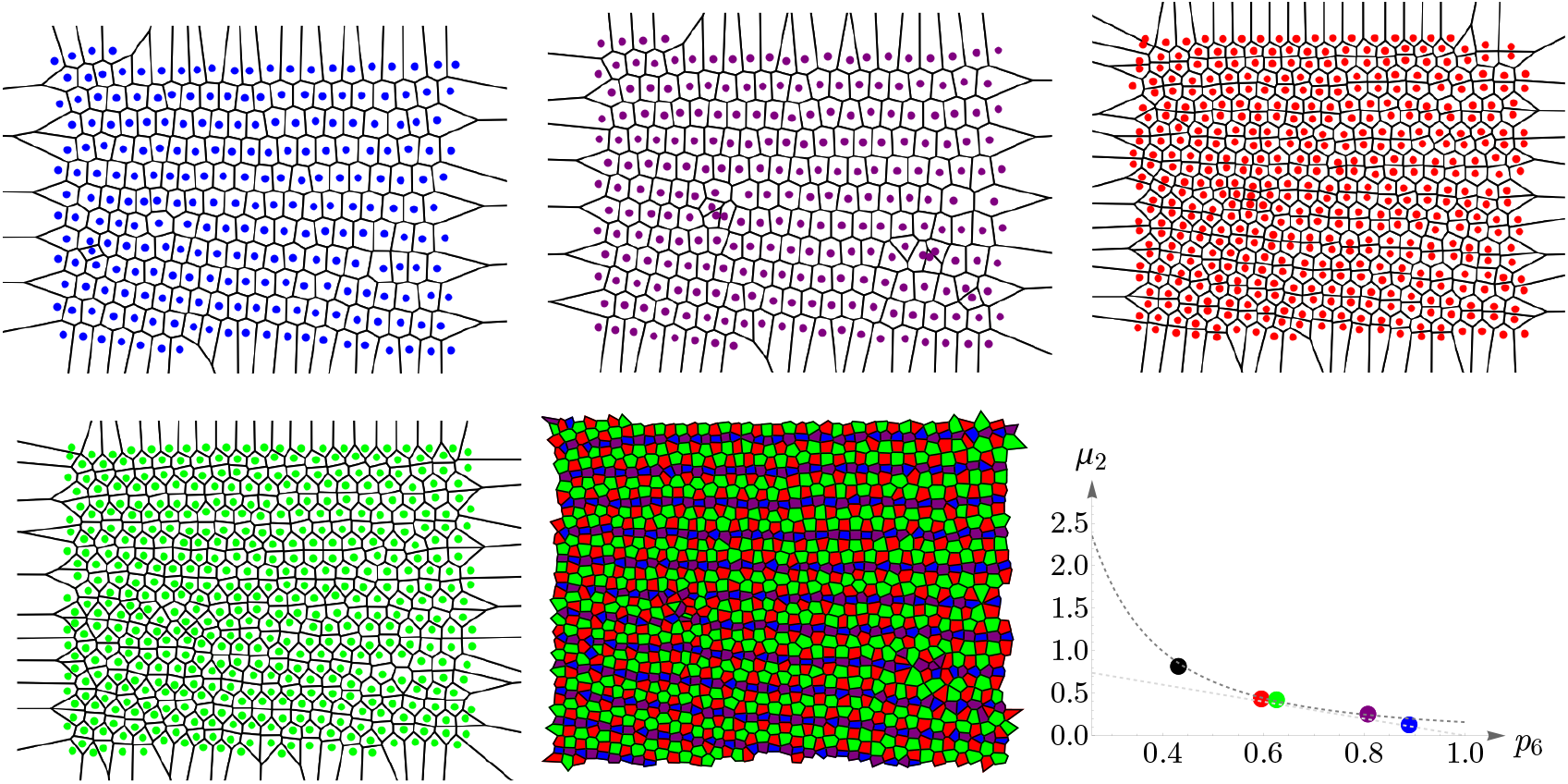
Voronoi tessellations of the zebrafish retinal field in Figure 9 and the corresponding cones’ behavior concerning Lemaître’s law. Each point in the plot corresponds to its respective cone subtype and the black point to the whole retinal field.

**Figure 24.**
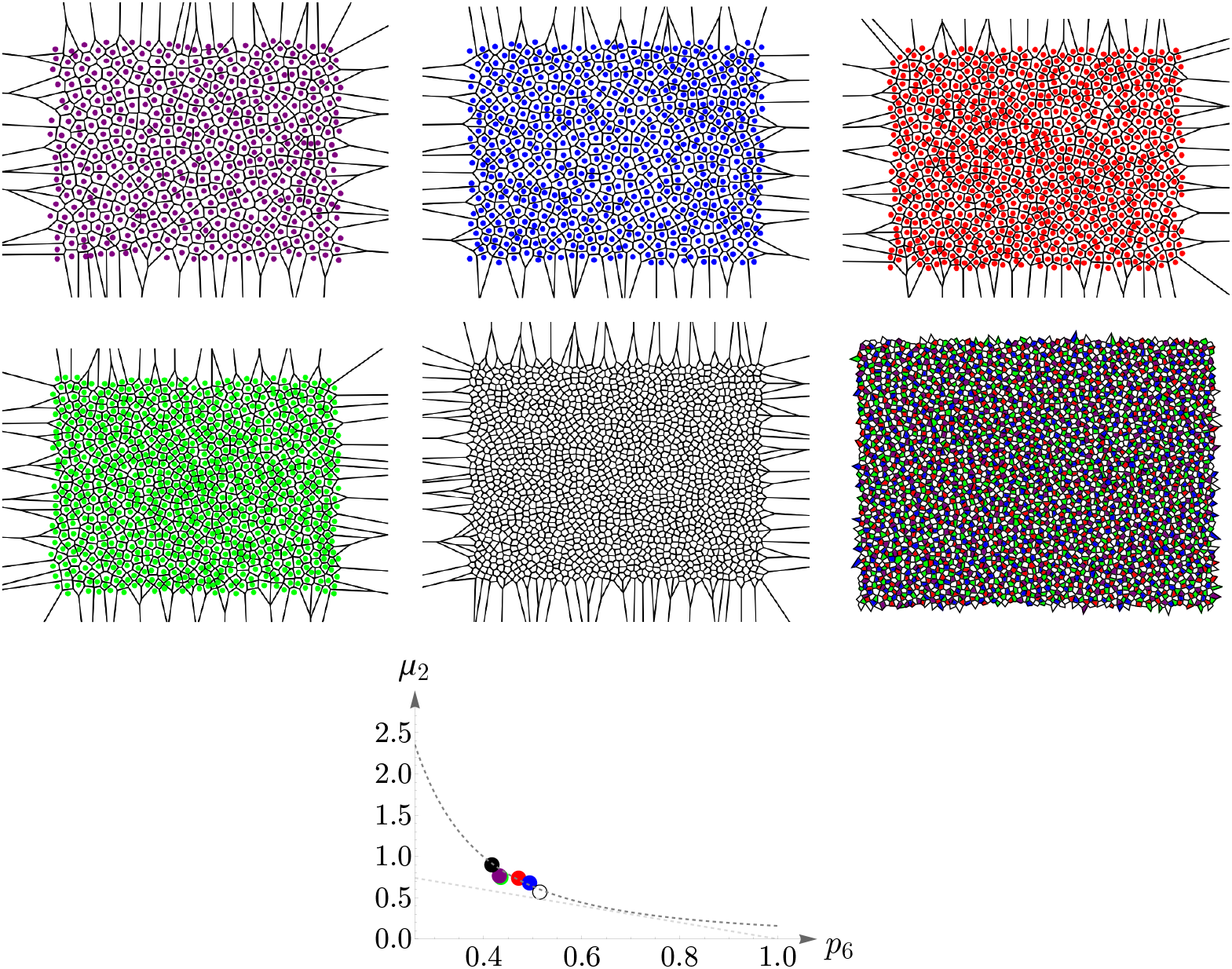
Voronoi tessellations of the chicken retinal field in Figure 10 and the corresponding cones’ behavior concerning Lemaître’s law. In the plot, the white point corresponds to double cones, and the black to the whole retinal field.

## 5. Concluding Remarks

In this work, we have applied the principle of maximum entropy to explain various forms of retinal cone mosaics in vertebrate eyes and established a parameter called retinal temperature or coldness, which is conserved throughout different species as diverse as rodent, dog, monkey, human, fish, and bird, regardless of the details of the underlying mechanisms, or physical and biological forces. This approach has enabled us to predict the frequency of the appearance of cone cells only by tuning a single parameter. The only constraint of the Lagrange problem stems from the repeatable nature of the experiments in biology.

Lemaître’s law which relates the fraction of hexagons to the width of the polygon distribution in numerous two-dimensional cellular networks in nature and is usually obtained by assuming an ad hoc constraint, here is derived as a special case of our formalism. We have shown that various networks, whether artificially generated or natural, obey this universal law.

Since we have considered a completely general constraint in the entropy maximization procedure, the approach of the current paper can be exploited to explain other patterns or processes in nature. In the case of failure, it implies that either additional information, which stems from the knowledge of the underlying mechanisms, needs to be considered, or the assumed information is incorrect. Indeed, this is one of the pitfalls of the maximum entropy approach as it is not falsifiable, and there are no criteria for its validity within itself [8,59].

Although in many cases, as in this paper, we can explain and predict the phenomena without knowing the details of the underlying dynamics, the principle of maximum entropy can still lead us to a better understanding of the involved mechanisms by validating the assumed information about the system.

## Institutional Review Board Statement

Not applicable.

## Informed Consent Statement

Not applicable.

## Data Availability Statement

We made secondary use of published data that can be found in the corresponding cited publications.

## Acknowledgments

The author is grateful to Carlo Beenakker and Iosif Pinelis for illuminating and inspiring discussions. He thanks Goethe University Frankfurt for providing financial and infrastructural support.

## Disclaimer/Publisher’s Note

The statements, opinions and data contained in all publications are solely those of the individual author(s) and contributor(s) and not of MDPI and/or the editor(s). MDPI and/or the editor(s) disclaim responsibility for any injury to people or property resulting from any ideas, methods, instructions or products referred to in the content.

